# Transcription starting inside human introns is widespread but mostly yields short-lived RNA

**DOI:** 10.64898/2026.11.24.768849

**Authors:** Michał W. Szcześniak, Elżbieta Wanowska

**Affiliations:** Institute of Human Biology and Evolution, Faculty of Biology, Adam Mickiewicz University in Poznań, Uniwersytetu Poznańskiego 6, 61-614 Poznań, Poland

**Author notes:** Corresponding author. Email addresses (MWS), (EW).

**Keywords:** Transcription initiation, Transcription start sites, Intronic transcription, PRO-cap, Long non-coding RNA, Nested genes, Nuclear exosome, Genome annotation

## Abstract

**Background:** Thousands of annotated long non-coding RNAs lie inside introns of protein-coding genes, on the host’s strand, but whether they are independent genes or pieces of host transcripts is usually unknown. An independent gene needs its own start site.

**Results:** We mapped transcription starts inside introns, using nascent capped RNA from two human cell lines. The approach detected the start sites of 94% of clearly expressed protein-coding genes nested in introns and confirmed 1,346 and 3,061 intronic start sites, almost all unannotated. Most are cell-type-specific, carrying active chromatin marks and motifs of the cell’s key transcription factors. Many initiate as strongly as the hosts’ alternative promoters but rarely sustain transcription: after a synchronised restart of transcription, their nascent RNA fades within a few kilobases, their short transcripts are partly degraded by the nuclear exosome, and the sequence downstream rarely has the early splice-donor site that protects promoter transcripts. Per initiation event, they leave about as little stable RNA as start sites between genes, so host transcription does not explain their low output. Expressed intronic RNAs on the host’s strand more often lack their own start site than those on the opposite strand.

**Conclusions:** Transcription starting inside human introns is widespread but mostly yields short-lived RNA. Many intronic long non-coding RNAs on the host’s strand are therefore likely to be pieces of host transcripts rather than independent genes, which matters for their annotation. All intronic start sites in both cell lines, with every measure used here, are provided as a resource.

## Background

Catalogues of human genes contain thousands of long non-coding RNAs. Among them is a class that lies inside the introns of protein-coding genes (host genes), on the same strand as the host [1]; the current GENCODE annotation lists more than two thousand such genes [2] (see Results). An RNA in such a position is hard to interpret. It may be an independent gene with its own promoter, as genes often lie entirely within another gene (nested genes): about a sixth of human genes contain at least one nested gene, most often within a single intron [3, 4]. It may also be a fragment of the host gene’s own transcript, such as a stable piece of an excised intron or a product of processing of the host’s precursor RNA (pre-mRNA), in which case it has no start site of its own. Expression data alone cannot tell these possibilities apart, because RNA from the host and RNA from an intronic promoter cover the same sequence on the same strand. What distinguishes an independent gene is that RNA polymerase II starts transcription at its 5′ end. Telling the two apart is a prerequisite for understanding the biological roles of intronic RNAs.

Transcription initiation in the human genome is pervasive. Upstream of active promoters, RNA polymerase II makes short, polyadenylated and unstable RNAs called promoter upstream transcripts (PROMPTs). They arise mostly from initiation in the antisense direction, away from the gene [5], and are degraded by the nuclear exosome, the main RNA-degrading complex of the nucleus [6]. Enhancers also initiate divergent transcription of short, unstable RNAs [7–9]. Whether initiation produces a stable RNA depends largely on the sequence downstream of the start. Transcribing towards the gene, the polymerase soon meets a splice-donor site (the start of an intron) and only later a polyadenylation signal, a sequence that triggers cleavage of the RNA and the end of transcription. In the opposite, PROMPT direction, polyadenylation signals are frequent; they are used, and the RNA is then degraded [5, 10]. Splice-donor sites bound by U1 small nuclear ribonucleoprotein protect the new RNA from premature cleavage and polyadenylation [11]. Many long non-coding RNAs between genes are also inefficiently processed and largely degraded in the nucleus [12].

Inside a transcribed gene, further factors could act. Transcription of the host gene could interfere with initiation inside it: in simpler systems such as bacteria, a polymerase passing through a promoter can dislodge the initiation machinery there or block its access, and the large introns of higher eukaryotes, which hold nested genes, offer ample opportunity for such interference [13]. In mouse embryonic stem cells, DNA methylation deposited by Dnmt3b, which the transcription-coupled mark H3K36me3 recruits to gene bodies, protects them from cryptic initiation [14]. Intragenic enhancers, conversely, can act as alternative promoters that produce spliced, polyadenylated RNAs [15], and their transcription can attenuate that of the host gene [16]. Transcription from a nested gene can even make transcription of its host end early. At the imprinted H13/Mcts2 locus in mouse, where expression depends on the parent of origin, H13 transcripts end inside the intron on the allele on which the nested Mcts2 gene is active [17], and silencing Mcts2 transcription lets H13 transcription reach its downstream ends [18]. For the host’s own strand, four questions have not been addressed genome-wide: how often transcription starts inside human introns, what kinds of elements these start sites are, whether, and how often, they give rise to stable RNA, and whether the host gene holds them back.

Here we approached annotated intronic RNAs from the other side: rather than starting from the RNAs themselves and their annotated 5′ ends, we mapped where transcription initiates inside introns, whether or not an RNA is annotated there. Using public PRO-cap data from two unrelated human cell lines, we mapped capped start sites inside introns and validated the approach on protein-coding genes known to be nested in introns. To follow what happens after initiation, we used nascent RNA after release from the transcription inhibitor DRB and RNA-seq from exosome knockouts, and we also analysed chromatin marks, conservation and sequence. We find thousands of intronic start sites, most of them cell-type-specific regulatory elements. Many initiate as strongly as the host genes’ internal promoters (annotated alternative promoters inside introns) but rarely sustain transcription. As a group, they have the sequence arrangement that protects promoter transcripts, a splice-donor site ahead of the first polyadenylation signal, barely more often than random intronic positions. Start sites between genes, which have no host, make about as little stable RNA per initiation event, so host transcription does not explain the low output of intronic start sites. Many annotated intronic RNAs that are expressed have no start site of their own and are likely pieces of the host’s transcript.

## Results

### Mapping transcription start sites inside introns

To find out which intronic RNAs could be genes in their own right, we did not start from the annotated RNAs. Instead, we asked where RNA polymerase II actually starts transcription inside introns, because an independent gene requires a start site of its own (Fig. 1a).

**Figure 1.**
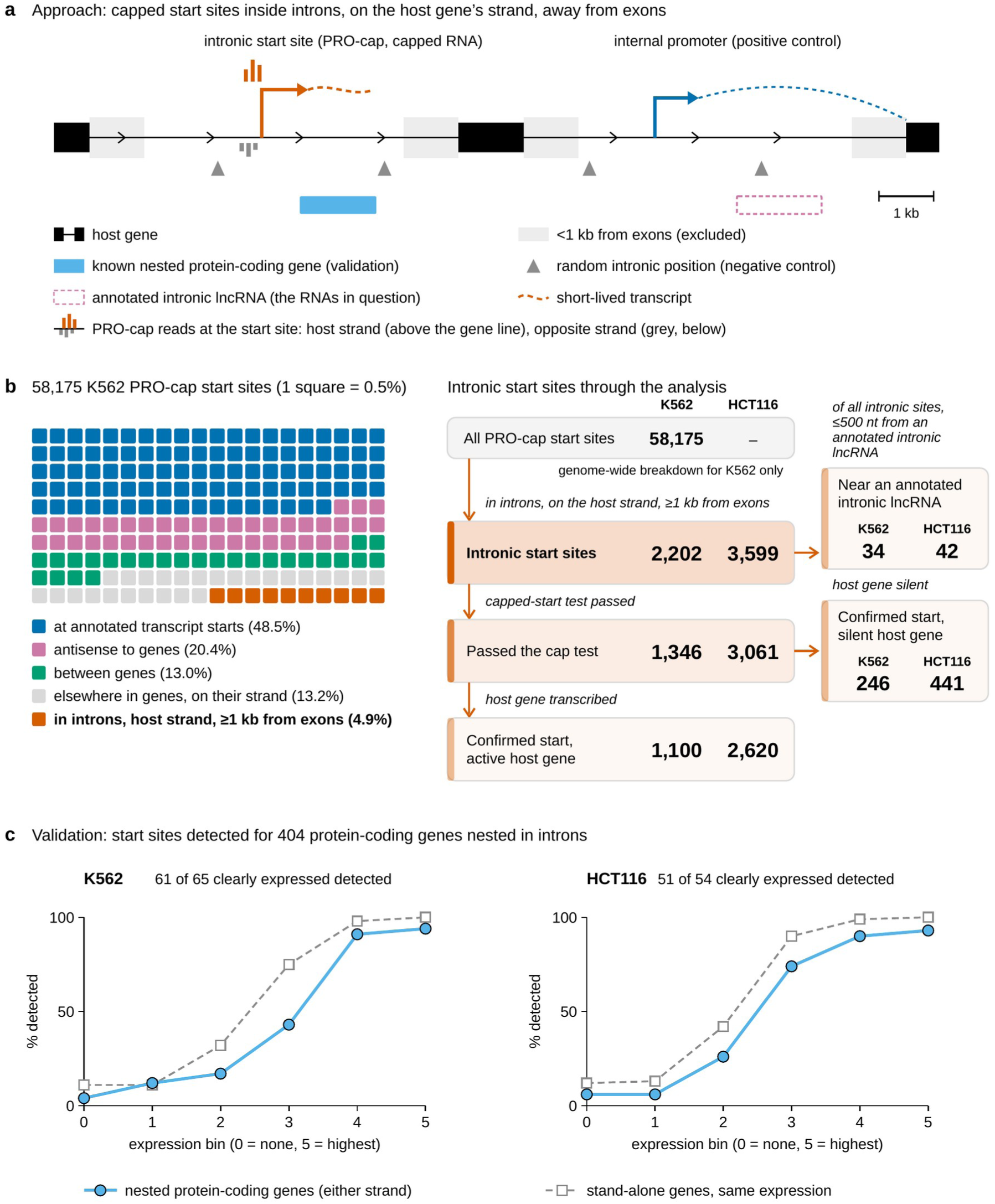
Transcription start sites inside introns. (a) The approach: capped start sites inside introns, on the host gene’s strand and at least 1 kb from its exons (light grey zones; bars above and below the gene line show PRO-cap reads on the host strand and on the opposite strand), with internal promoters as positive and random intronic positions as negative controls, and protein-coding genes nested in introns for validation. (b) Genomic location of all PRO-cap start sites in K562 (each square 0.5%), and the intronic start sites followed through the analysis in both cell lines (K562 / HCT116), including those in silent host genes. (c) Start sites detected for protein-coding genes nested in introns of other genes, compared with stand-alone genes of the same expression, by expression bin of the gene itself.

We used PRO-cap, which maps the capped 5′ ends of newly made RNAs and therefore marks transcription start sites at nucleotide resolution [19, 20], in two unrelated human cell lines from ENCODE: the leukaemia line K562, for which a matched library of uncapped RNA serves as background, and the colon cancer line HCT116. For HCT116 we used untreated cells of two lines engineered for rapid degradation of CTCF or RAD21 (degron lines); the CTCF line, with deeper data, served for the main analyses and the RAD21 line as an independent replicate. Because RNA polymerase II caps new RNA right after initiation, a capped 5′ end marks a start site. A site counted as a confirmed start when it passed a cap test: enough capped reads, clearly more than in its flanks and, in K562, more than in the uncapped library (Methods). We considered start sites inside introns of protein-coding genes, on the host gene’s strand and at least 1 kb from its exons (Fig. 1b), and called a host gene active when its own start carried at least five capped reads. Readthrough genes, which GENCODE annotates across two or more neighbouring genes, were excluded as hosts, because they would turn the promoter of the downstream gene into an apparent intronic start site [2].

Before interpreting new start sites, we tested whether the approach detects known ones: protein-coding genes nested inside an intron of another protein-coding gene, whose status as genes is not in doubt (404 in GENCODE). Of those clearly expressed in each cell line, 94% had a detected start site, about as often as stand-alone genes of the same expression (Fig. 1c). Every nested gene that is well established (with a MANE Select transcript) and broadly expressed across tissues was detected (50 of 50 in K562 and 42 of 42 in HCT116; Methods), among them NSMCE3 inside ENTREP2, CHMP1B inside GNAL and JUNB inside HOOK2. The few clearly expressed nested genes without a detected start (4 in K562 and 3 in HCT116) were mostly recently annotated genes without a name or genes in complex loci, such as OR51B4 in the β-globin cluster and HOXA5 in the HOXA cluster. The cap test also confirmed about a tenth of the host genes’ annotated alternative promoters inside introns. We call these internal promoters and use them as positive controls; below, the term means those that passed the cap test, unless all annotated ones are meant. By contrast, fewer than 1% of random positions in the same introns passed (Additional file 1: Fig. S1). In HCT116, 91% of the confirmed sites passed the cap test again in the independent RAD21 line, including 79% of the third with the fewest PRO-cap reads, which are the most likely to be missed (Additional file 1: Fig. S2).

With this validation in hand, we mapped intronic start sites genome-wide. In K562, 4.9% of the 58,175 PRO-cap start sites lay in introns of protein-coding genes, on the gene’s strand and at least 1 kb from exons (Fig. 1b). After merging nearby summits and applying stricter host rules (Methods), we obtained 2,202 intronic start sites in K562 and 3,599 in HCT116 (Additional file 2: Table S1). Of these, 1,346 and 3,061 passed the cap test (a larger share in HCT116, where no uncapped control was available and the test was therefore less strict): 1,100 and 2,620 in active host genes, which form the main set analysed below, and 246 and 441 in silent ones (Fig. 1b). Almost none coincided with annotated starts: of all 2,202 K562 and 3,599 HCT116 sites, 96% and 97% lay more than 500 nt from any annotated transcript start on the same strand, and only 34 of the K562 sites lay within 500 nt of an annotated intronic long non-coding RNA, so current annotations capture few of the places where transcription starts inside introns.

### Most intronic start sites are cell-type-specific regulatory elements

Intronic start sites carry the chromatin marks of active regulatory elements (Fig. 2a). Nearly 90% of the confirmed sites in active genes had a peak of H3K27ac, the histone mark of active promoters and enhancers, within 1 kb, against 12–17% of random intronic positions. In K562, about half also carried the promoter mark H3K4me3, as expected of a mixture of promoter-like and enhancer-like elements; in HCT116 most did, but the histone data of the two cell lines differ in depth and quality, which changes how many peaks are called, so this share is not comparable between them (Fig. 2 legend). Intronic start sites were also more conserved in evolution than their surroundings: 7–9% had a mean phyloP score above 1 (a measure of how slowly a position has changed across 100 vertebrates), against 1–2% of random intronic positions and 22–23% of host promoters.

**Figure 2.**
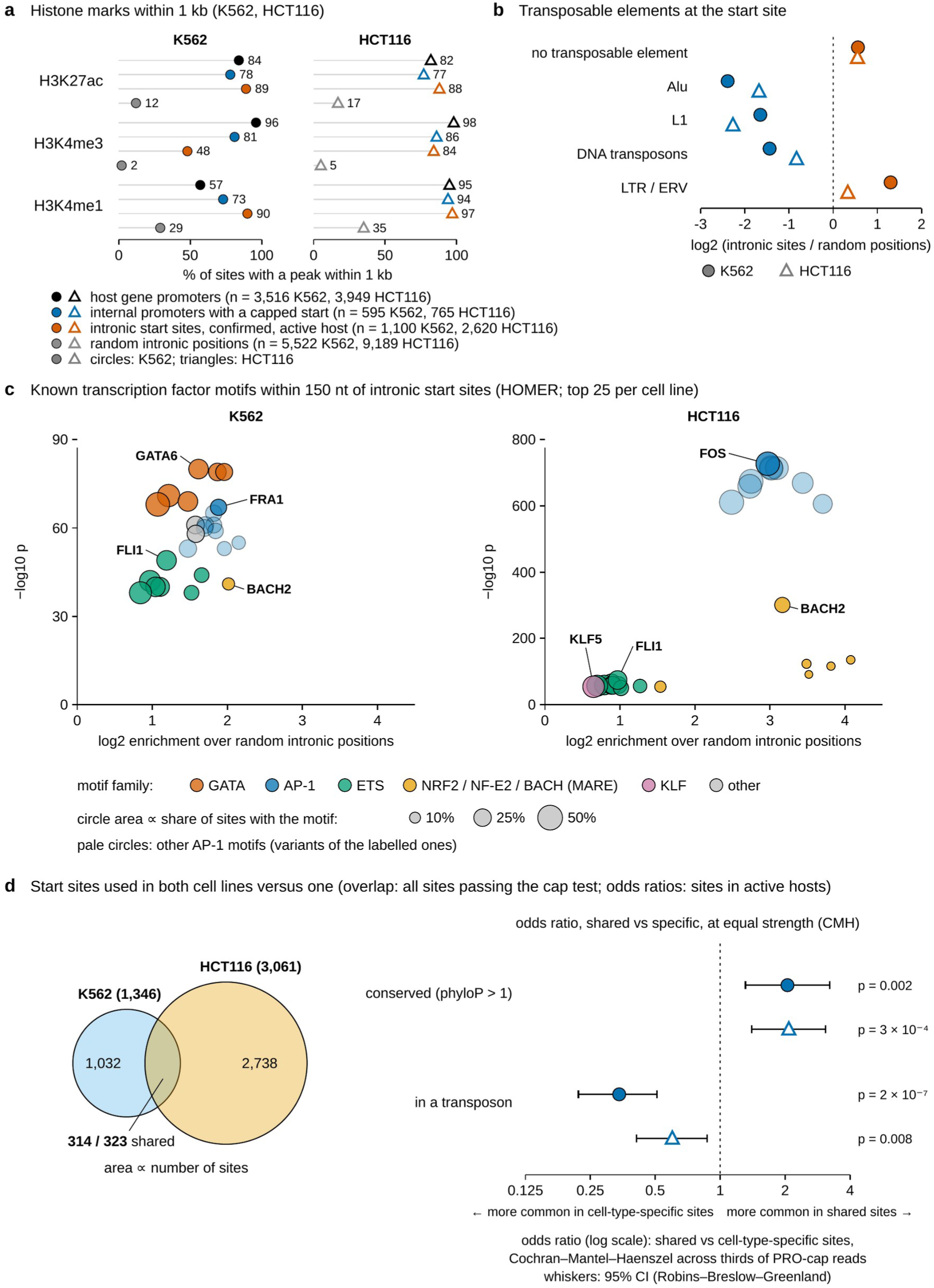
Intronic start sites are cell-type-specific regulatory elements. (a) Histone marks within 1 kb of host gene promoters, internal promoters with a capped start, intronic start sites in active hosts that passed the cap test, and random positions in the same introns; circles K562, triangles HCT116. Histone peak sets differ between the cell lines in depth and quality (for example, 95% of host promoters carry an H3K4me1 peak in HCT116, against 57% in K562), so shares are best compared within a cell line. (b) Transposable element classes at intronic start sites relative to random intronic positions (log2 ratio of shares; circles K562, triangles HCT116). Sites in any transposable element: 26% (K562) and 23% (HCT116), against 49% and 47% of random positions; LTR/ERV elements: 10.6% against 4.3% (p = 1 × 10⁻¹⁴) and 5.4% against 4.3% (p = 0.02); LTR12C: 4 and 8 sites. (c) Known transcription factor motifs (HOMER, top 25 per cell line) within 150 nt of intronic start sites: enrichment against significance, coloured by motif family; circle area shows the share of sites with the motif; pale circles are further variants of the AP-1 motif; p values from the HOMER binomial test (top-ranked motifs: GATA, p = 10⁻⁸⁰, in K562; FOS, p = 10⁻⁷²⁷, in HCT116). (d) Start sites used in both cell lines versus one: overlap of all sites that passed the cap test (314 K562 and 323 HCT116 sites lie within 200 nt of a site in the other cell line), and odds ratios (log scale, with 95% confidence intervals) for conservation and transposon origin of shared against cell-type-specific sites in active hosts at equal initiation strength (Cochran–Mantel–Haenszel test, which combines the three thirds of PRO-cap reads); an odds ratio above 1 means that the feature is more common among shared sites, below 1 that it is more common among cell-type-specific sites.

Transposable elements carry ready-made promoters, and these can be reactivated in cancer cells [21]. Intronic start sites were nevertheless depleted of them: about a quarter lay in a transposable element, against nearly half of random intronic positions (Fig. 2b). Alu, L1 and DNA transposons were all depleted, and only the long terminal repeats (LTRs) of endogenous retroviruses were enriched, 2.5-fold in K562 and 1.25-fold in HCT116. They included a few copies of LTR12C, a member of the ERV9/LTR12 family whose copies serve as cryptic promoters in cancer cells treated with inhibitors of DNA methylation or histone deacetylation [22]. Three quarters of intronic start sites thus lie outside annotated transposable elements.

The sequences around intronic start sites carried the binding motifs of each cell type’s key transcription factors (Fig. 2c). In K562, GATA motifs ranked first: those of GATA1 and GATA2 were present at a quarter of sites, against 6–7% of random intronic positions. Both are haematopoietic factors, and GATA1 is regarded as the master regulator of red blood cell development [23]. They were followed by motifs of AP-1 and ETS factors and the Maf recognition element (MARE) bound by NF-E2 and BACH factors. In HCT116, motifs of AP-1, a family of FOS and JUN factors, dominated, with a FOS motif at 45% of sites against 6% of random positions, followed by the MARE motif, ETS motifs and the motif of KLF5, a transcription factor of dividing cells of the intestinal epithelium [24]. Together, these features indicate that most intronic start sites are regulatory elements of each cell’s own transcriptional programme that lie inside genes and initiate transcription.

Most intronic start sites were specific to one cell type (Fig. 2d): only 23% of the K562 sites that passed the cap test had a counterpart within 200 nt in HCT116, and only 11% of the HCT116 sites one in K562. Weak sites are more easily missed in one line, but even among the strongest third, only a third of the K562 sites and a fifth of the HCT116 sites were found in the other cell line. Nor did this specificity simply follow host activity: 60% of the K562-specific sites lay in genes that are also active in HCT116. Compared with cell-type-specific sites of the same initiation strength, the sites used in both cell lines had about twice the odds of being conserved and lower odds of lying in a transposable element (Fig. 2d), but they did not give stable RNA more often (odds ratios 1.01 and 1.54, p = 0.95 and 0.06; stable RNA is measured as described in the next section).

### Intronic start sites initiate transcription but rarely sustain it

Initiation alone does not make a gene: most genes also produce RNA that persists in the cell, although some long non-coding RNAs are unstable and may act through their transcription rather than their product. For each site, we therefore asked whether stable RNA is made from it, that is, whether total RNA-seq coverage on the site’s strand was higher in the 3 kb after the start than in the 3 kb before it, beyond the 95th percentile of random intronic positions (Methods). We compared intronic start sites with internal promoters of similar initiation strength, divided into thirds by the number of PRO-cap reads at the site. In active hosts, sites closer than 3 kb to the next exon of the host gene were left out, because the window after the site then contains that exon, whose coverage by spliced host mRNA mimics such a rise (Additional file 1: Fig. S9a). Even in the strongest third, a rise was far less common at intronic start sites (23% in K562 and 14% in HCT116) than at internal promoters of the same strength (75% and 67%; Fig. 3a). Intronic start sites had somewhat fewer PRO-cap reads than internal promoters on average (Fig. 3 legend), but the two distributions overlapped widely, and more than a quarter of intronic start sites fell into the strongest third, that is, initiated as strongly as the stronger internal promoters. Even at equal initiation strength, intronic start sites far less often produced stable RNA, so weaker initiation does not explain their low output.

**Figure 3.**
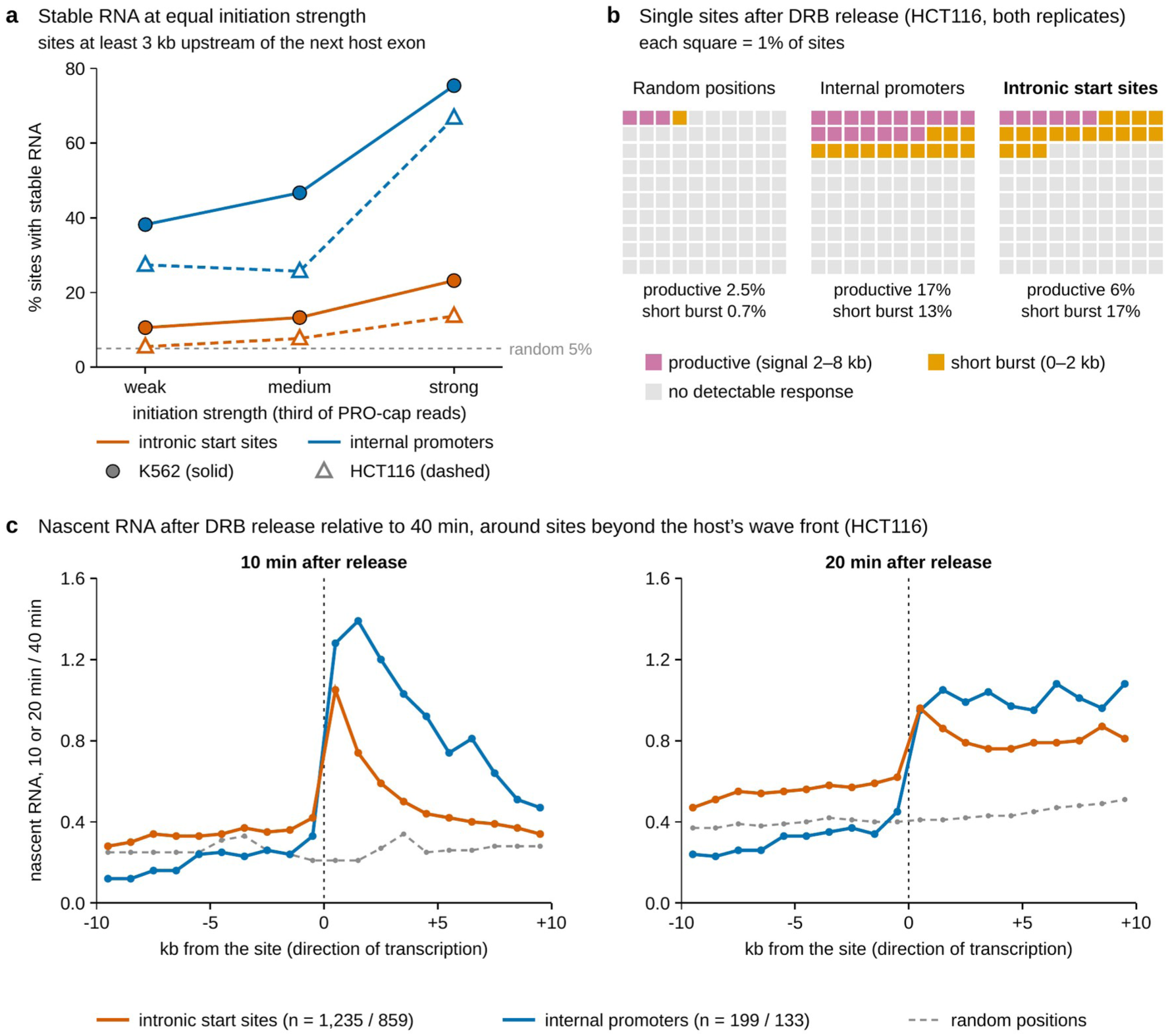
Intronic start sites initiate transcription but rarely sustain it. (a) Share of intronic start sites and internal promoters with a rise in stable RNA above the 95th percentile of random intronic positions, by third of initiation strength; only sites at least 3 kb upstream of the next exon of the host gene, because closer to an exon the host’s mRNA can mimic a rise (Methods); K562 solid, HCT116 dashed; horizontal dotted line, random positions (5% by construction, because the threshold is their 95th percentile). Median PRO-cap reads per site: intronic start sites 42 (K562) and 49 (HCT116), internal promoters 56 and 114; 300 of 1,100 (K562) and 723 of 2,620 (HCT116) intronic start sites fall in the strongest third. (b) Single sites classified from both replicates of the HCT116 time course: intronic start sites in active hosts (n = 1,338), internal promoters (n = 249) and random intronic positions (n = 4,353); productive (signal 2–8 kb after the site), short burst (0–2 kb only) or no detectable response. (c) Nascent RNA 10 and 20 minutes after DRB release relative to 40 minutes, around sites beyond the front of the host gene’s own wave (HCT116; at 10 minutes 1,235 intronic start sites in active hosts and 199 internal promoters, at 20 minutes 859 and 133).

To see what happens after initiation, we followed newly made RNA after a synchronised restart of transcription. The drug DRB, a reversible inhibitor of the kinase CDK9, holds RNA polymerase near promoters; after it is washed out, a wave of polymerases leaves active promoters at the same time [25, 26]. In HCT116, newly made RNA was labelled and sequenced 10 to 40 minutes after release (DRB/TTchem-seq) [27]. Ten minutes after release, the host genes’ own polymerases had reached 12–20 kb into the genes; we call this zone the wave front (Additional file 1: Fig. S3). We therefore tested only intronic start sites well beyond the wave front (Methods), so that new RNA downstream of them could not come from the host promoter. As a group, these 1,191 sites produced a burst of new RNA, whereas random intronic positions produced none (p = 3 × 10⁻⁵⁰, one-sided Mann– Whitney test). Similar shares of intronic start sites and internal promoters responded, that is, exceeded the 95th percentile of random positions (15% and 20%), but the burst was much smaller at intronic start sites (see the average profiles below). The response did not come from internal promoters farther upstream, whose polymerases could reach an intronic site within 10 minutes: without the 38 sites that had an active internal promoter up to 20 kb upstream, the share was unchanged (Methods). A lower-depth DRB dataset from K562 was not sensitive enough to detect this small burst reliably (Additional file 1: Fig. S4).

The two replicates of the HCT116 time course also allowed each site to be assigned to a response class (Fig. 3b): no detectable response, a short burst confined to the first 2 kb, or transcription extending further, which we call productive. Of the 1,338 intronic start sites in active genes that could be classified (enough reads and beyond the host’s wave front; Methods), most (77%) showed no detectable response, 17% a short burst and 6% productive transcription. Only 0.7% of random positions were classed as short, so about one short call in 25 is expected to be false, but 2.5% of random positions were classed as productive, so about half of the productive calls may be false. Internal promoters, by contrast, were classed as productive more often than as short. The test compares the signal 10 minutes after release with that at 40 minutes, when the host’s own polymerases have reached the site; in strongly transcribed hosts this reference is high, so a far-reaching burst from an intronic site can go unnoticed, and the share of productive sites may be underestimated.

On average, the burst was short (Fig. 3c). New RNA from intronic start sites peaked in the first kilobase, and 4–5 kb downstream less than a fifth of this excess over the level before the site remained, whereas at internal promoters more than half remained. After 20 minutes, a small excess was present across the whole 10 kb downstream of intronic start sites, presumably carried by the minority of polymerases that kept going.

Short RNAs produced by early-terminating transcription, such as PROMPTs, are degraded in the nucleus by the RNA exosome [5, 6], with the help of the RNA helicase MTREX, part of the nuclear exosome targeting (NEXT) complex [28]. ENCODE provides RNA-seq from K562 cells in which MTREX, EXOSC10 or EXOSC5 was knocked out. Judged by PROMPTs, classic exosome targets, only the MTREX knockout worked, and only partially (Additional file 1: Fig. S5). Compared with control cells, RNA downstream of intronic start sites rose in these cells more often than at random positions: 12% of intronic start sites in active hosts exceeded a threshold passed by 5% of random positions (Fig. 4a). The rise came mainly from promoter-like sites, those with H3K4me3, and was strongest in silent hosts (41% of sites), where no host transcript covers the intron and dilutes the signal. RNA from internal promoters, which is mostly stable and spliced, did not rise significantly more often than at random positions. Because the knockout was partial and the libraries captured only polyadenylated RNA, these effects are probably underestimated. Intronic start sites therefore initiate transcription, but most of their polymerases stop within a few kilobases, and at least part of the short RNA they leave is degraded by the nuclear exosome.

**Figure 4.**
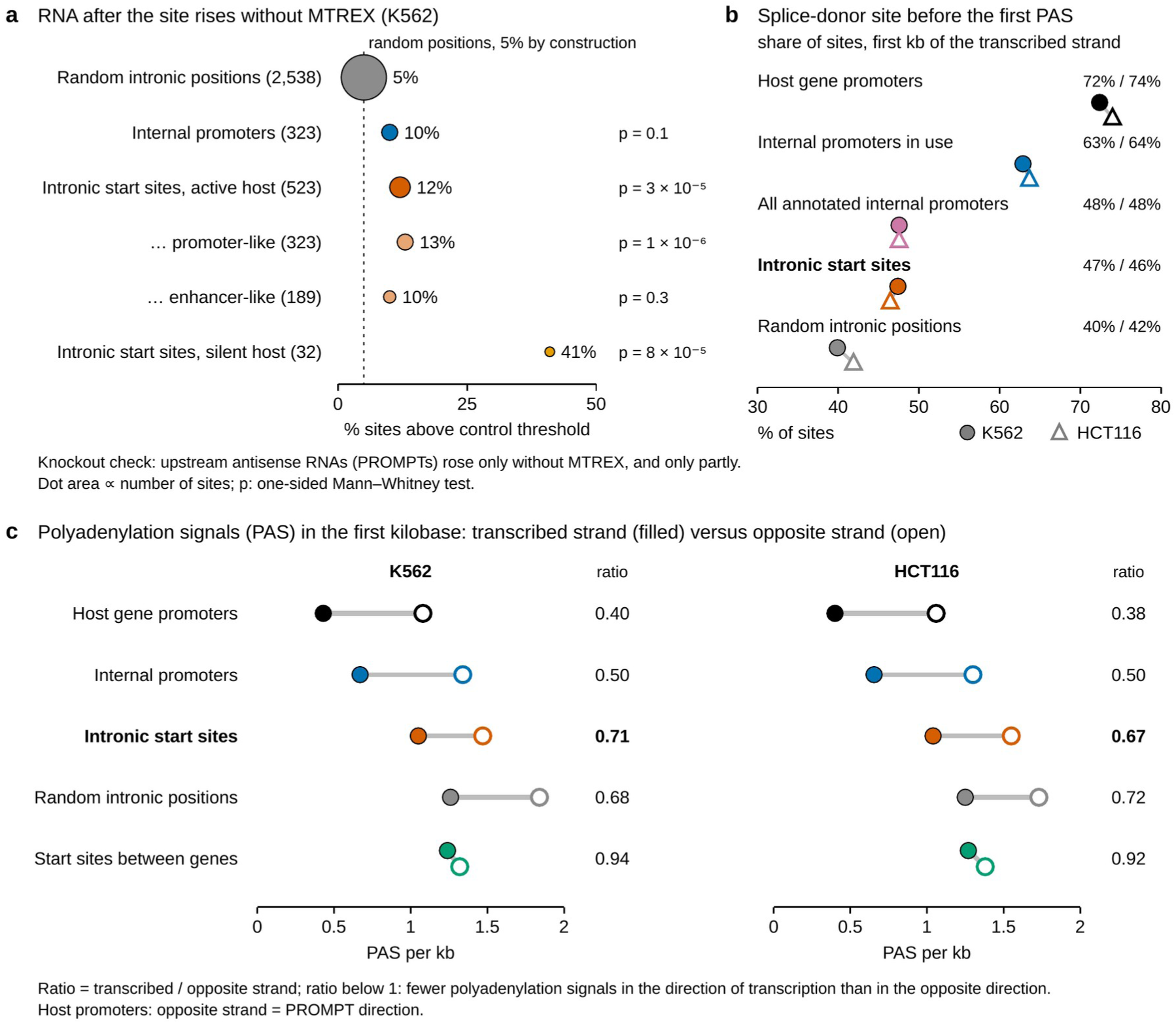
Short RNAs are partly degraded, and the sequence downstream resembles the intron. (a) Share of sites whose downstream RNA rises in K562 cells lacking MTREX, relative to control cells, above the level reached by 95% of random intronic positions; dot area is proportional to the number of sites, given in brackets (the smallest dot is enlarged to stay visible); one-sided Mann–Whitney tests against random positions. (b) Share of sites at which a splice-donor motif precedes the first polyadenylation signal in the first kilobase on the transcribed strand, or no signal is present, for host gene promoters, internal promoters in use, all annotated internal promoters, intronic start sites in active hosts and random intronic positions. (c) Polyadenylation signals (PAS) per kb in the first kilobase on the transcribed strand (filled) and on the opposite strand in the opposite direction (open), with their ratio, for the groups in (b) except all annotated internal promoters, and for start sites between genes. Within intronic start sites, those with the protective arrangement gave stable RNA in 17.0% against 13.4% of cases in K562 (p = 0.14) and 9.5% against 7.8% in HCT116 (p = 0.17; Fisher’s exact test).

### Downstream of intronic start sites, the sequence resembles the intron rather than a promoter

We next asked whether the sequence downstream of the start differs between intronic start sites and promoters. At promoters of genes, this sequence favours productive transcription in the direction of the gene [5, 10], so if intronic start sites were promoters of independent genes, theirs should do the same. As outlined in the Introduction, transcription at promoters soon meets a splice-donor site, which protects the new RNA from early cleavage, and only later a polyadenylation signal (the hexamer AATAAA or ATTAAA), which triggers cleavage and the end of transcription [10, 11, 29]. This protective arrangement, a splice-donor motif before the first polyadenylation signal in the first kilobase, or no signal at all, was present at about three quarters of host promoters and two thirds of internal promoters in use, but at fewer than half of intronic start sites, close to random intronic positions (Fig. 4b). Annotated internal promoters taken together, used or not, had it at 48%, as often as intronic start sites, so the arrangement distinguishes internal promoters that are used from those that are merely annotated. Within intronic start sites, it made little difference: sites with the arrangement gave stable RNA only slightly, and not significantly, more often in both cell lines.

Polyadenylation signals alone told the same story (Fig. 4c). We compared their number in the first kilobase on the transcribed strand with that on the opposite strand, read away from the site, as in the PROMPT direction at promoters. Host promoters and internal promoters carried half as many signals or fewer in the direction of transcription, intronic start sites only about a third fewer, like random positions in the same introns. This mild shortage is expected from base composition: the strand of an intron that is transcribed with the host is richer in G and T [30] and so carries fewer of the A-rich signals by chance. Start sites between genes showed almost no asymmetry, in line with the background-level frequencies of polyadenylation signals and splice-donor motifs reported downstream of enhancer start sites [9]. The sequence downstream of intronic start sites thus resembles the surrounding intron rather than that of promoters. Polyadenylation signals also mattered at single sites: sites without a detectable response after DRB release had more of them in the first kilobase than sites with a short or productive response (1.12 against 0.91 and 0.84 per kb; p = 0.007, Mann–Whitney test), which did not differ from each other.

### Host transcription does not explain the low output of intronic start sites

Besides the downstream sequence, a second possible explanation for the low output is location: polymerases passing through from the host gene might interfere with intronic start sites. RNA made at enhancers and other regulatory elements between genes is also short-lived and degraded by the exosome [9], so start sites between genes are the natural reference for asking whether lying inside a transcribed gene changes what intronic start sites produce. We compared them with start sites at least 5 kb from any protein-coding gene (4,669 in K562 and 10,318 in HCT116). All sites in this comparison passed the cap test, lay more than 500 nt from any annotated transcript start and were compared within thirds of initiation strength (Methods).

This comparison needs a measure that the host gene does not distort. In an active gene, the host’s precursor RNA already covers the intron before an intronic start site, so the ratio of coverage after the start to coverage before it is pulled towards 1 and makes intronic sites look several times weaker than sites between genes (Additional file 1: Fig. S6). We therefore used the difference in RNA-seq coverage after and before the start, divided by the number of PRO-cap reads at the site, as an estimate of the stable RNA made per initiation event (strictly, per PRO-cap read: PRO-cap mostly counts polymerases paused near the start, so a site where they stay paused longer gives more reads without starting more transcripts). By this measure, intronic start sites in active and in silent genes and start sites between genes all made very little stable RNA, whereas internal promoters made far more (Fig. 5a). Intronic start sites in active genes had more coverage after the start than before it at fewer sites than start sites between genes, clearly in K562 (44% against 56%) and slightly in HCT116 (43% against 46%), but at more sites than random positions in the same introns. The slightly lower share at intronic start sites may partly reflect the host’s precursor RNA, whose coverage tends to fall along an intron from its 5′ to its 3′ end, so that the coverage after any site is slightly lower than before it. Nascent RNA after DRB release gave a similar picture (Fig. 5b). Per PRO-cap read, intronic start sites in active genes produced more new RNA 2–8 kb downstream than start sites between genes, that is, more of their polymerases travelled that far, but about ten times less than internal promoters; intronic start sites in silent genes did not differ from start sites between genes. Being inside an active gene therefore does not explain why intronic start sites make so little stable RNA: start sites between genes, which have no host, make about as little. Whether host transcription reduces initiation itself cannot be judged from these sites alone (but see the strand comparison below).

**Figure 5.**
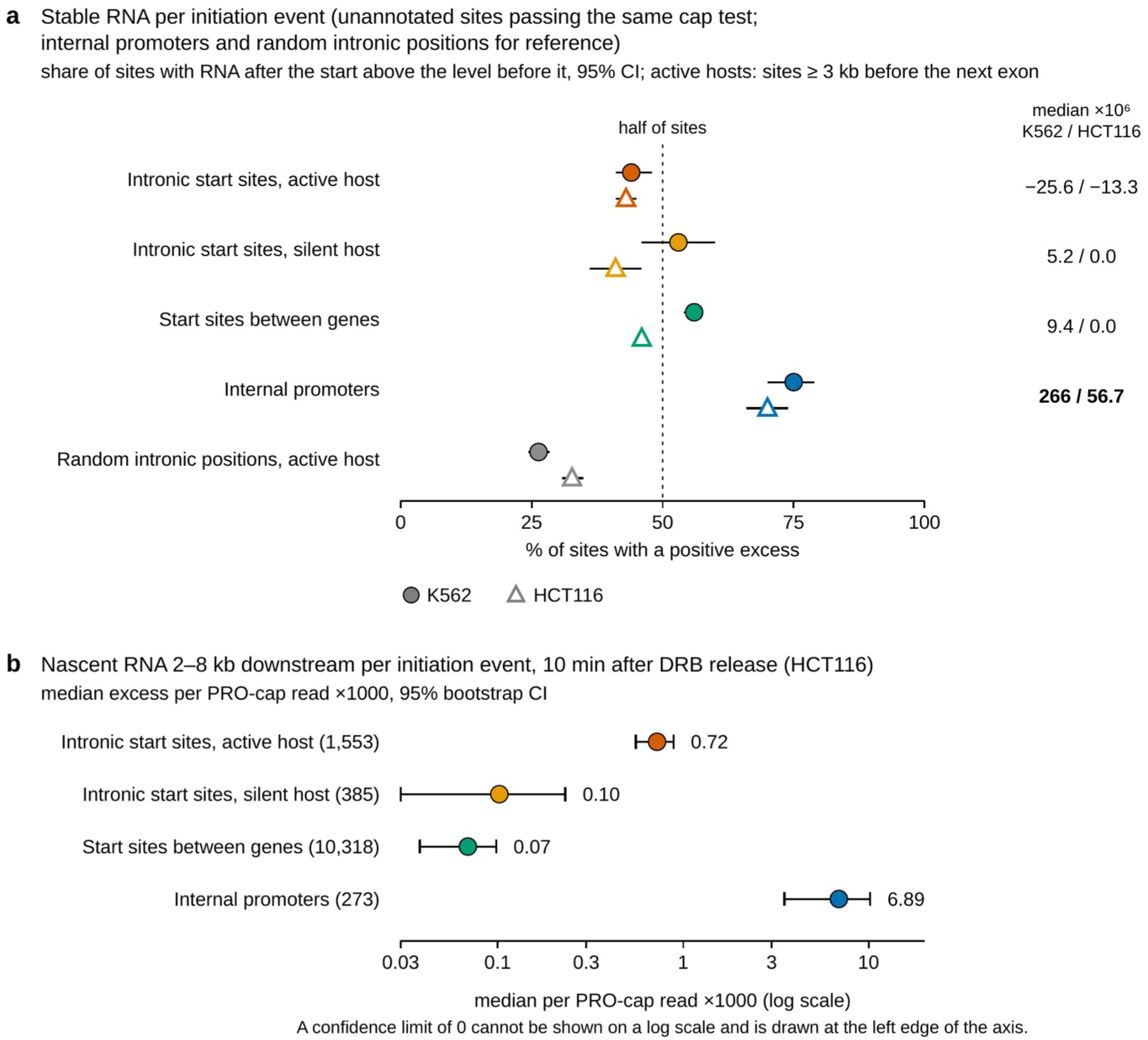
Host transcription does not explain the low output of intronic start sites. Unannotated start sites passing the same cap test, compared by RNA produced per initiation event (per PRO-cap read at the site); annotated internal promoters and, in (a), random intronic positions in active hosts are shown for reference. (a) Share of sites with more stable RNA after the start than before it, with 95% confidence intervals, and median excess of RNA-seq coverage per PRO-cap read (in units of 10⁻⁶); intronic start sites in active hosts, internal promoters and random positions only at least 3 kb upstream of the next exon of the host gene (Methods); random positions have no PRO-cap reads, so no median is shown for them. (b) Nascent RNA 2–8 kb downstream per PRO-cap read 10 minutes after DRB release (HCT116; 1,553 intronic start sites in active hosts, 385 in silent hosts, 10,318 start sites between genes and 273 internal promoters), median excess (in units of 10⁻³) with 95% bootstrap confidence interval, log scale.

Figure 6 shows these patterns at single loci: intronic start sites whose nascent RNA fades within a few kilobases (RAI1; Fig. 6a) or reaches further (ZMIZ1; Fig. 6b), in both cases without stable RNA. For comparison, Additional file 1: Fig. S10 shows a protein-coding gene nested in an intron (CHMP1B in GNAL) and an internal promoter whose RNA joins the host’s exons (ACIN1).

**Figure 6.**
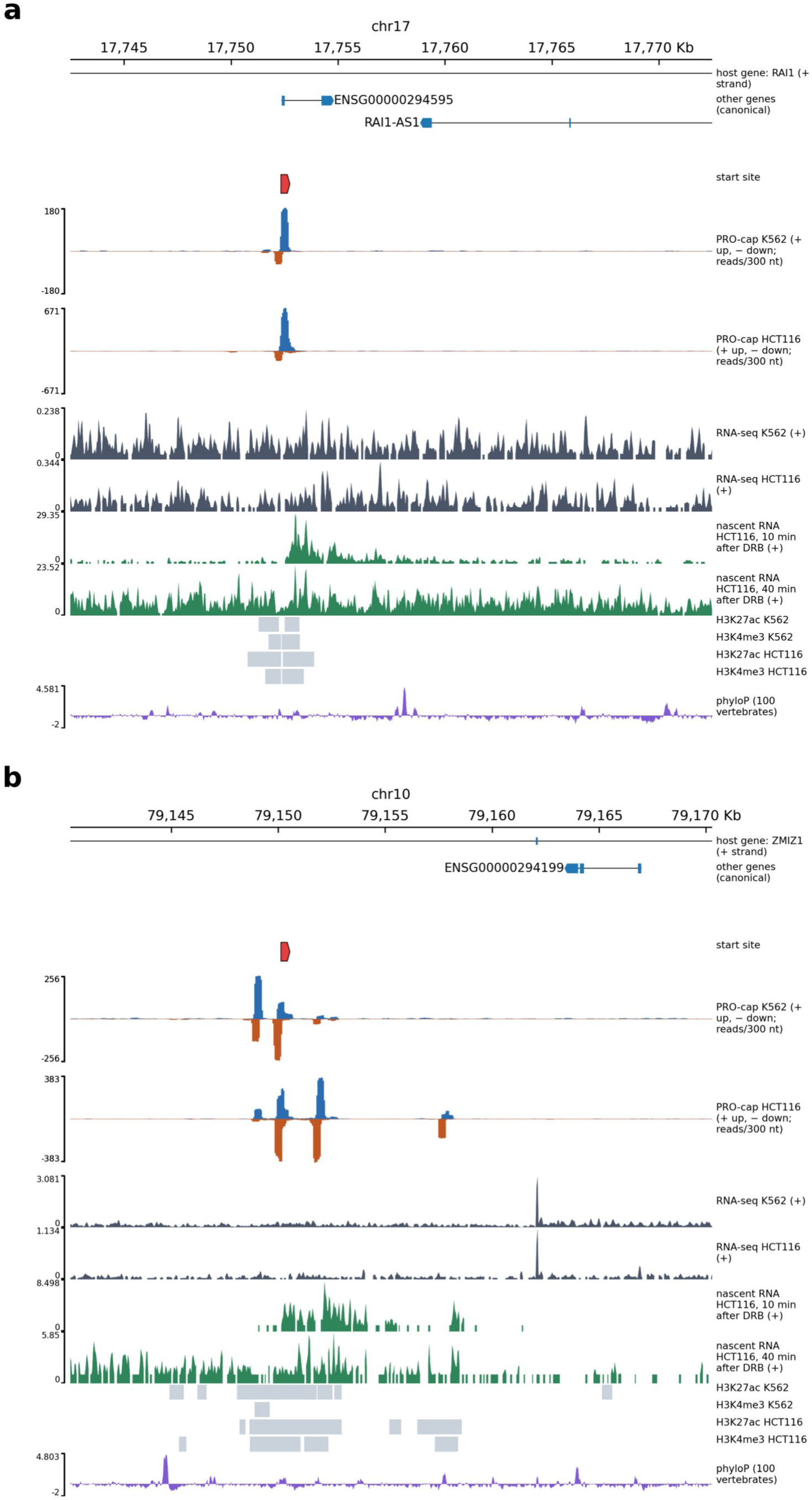
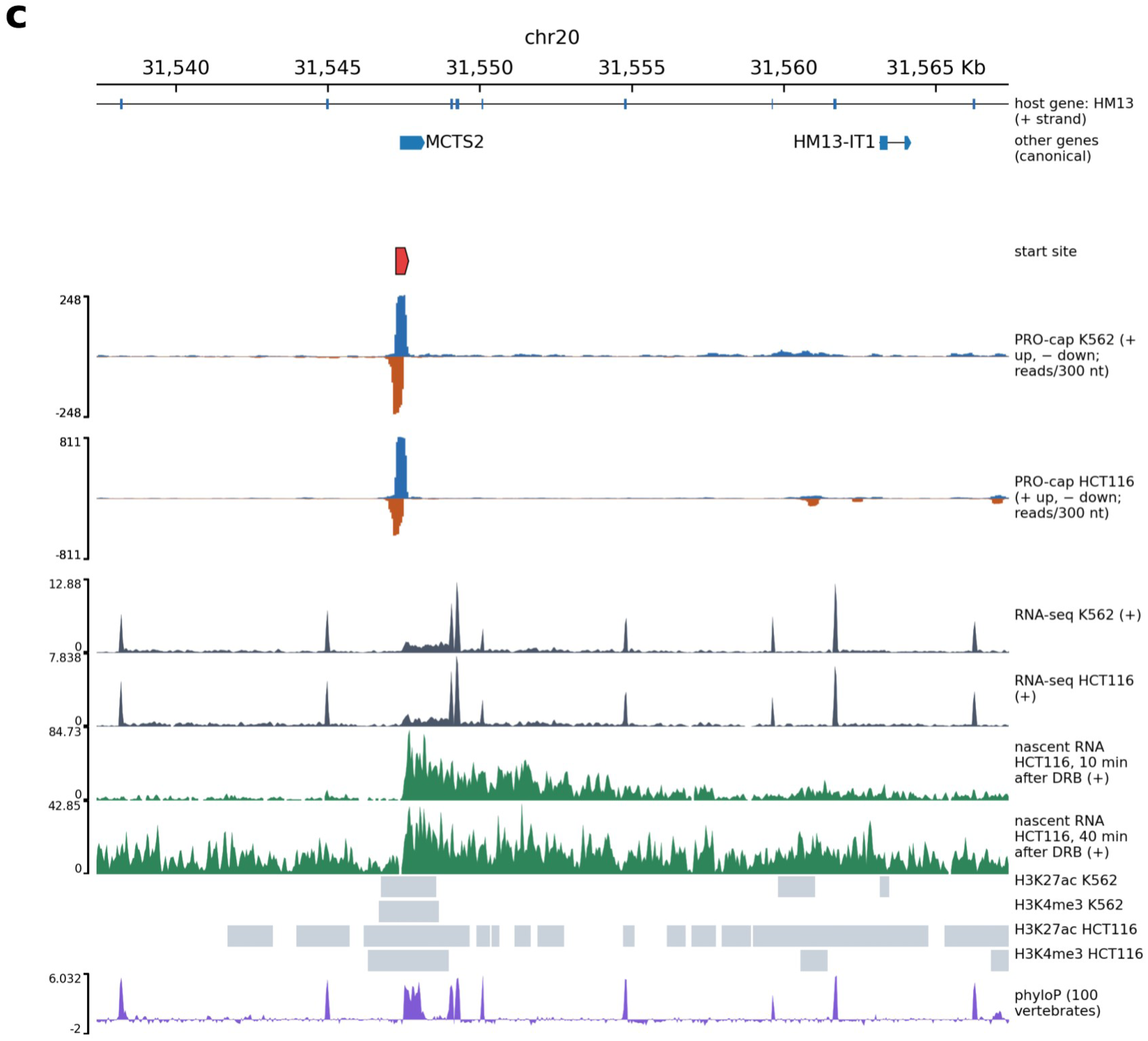
Examples at single loci. Each panel shows, from top: the host gene (with its strand) and other genes (canonical transcripts), the start site (red), PRO-cap on both strands in K562 and HCT116 (reads summed in 301-nt windows centred on each position, so that narrow start peaks remain visible at this scale; + strand up, − strand down), total RNA-seq on the site’s strand in both cell lines, nascent RNA 10 and 40 min after DRB release in HCT116, H3K27ac and H3K4me3 peaks in both cell lines, and conservation (phyloP). The pointed ends of genes and of the start-site mark show the direction of transcription. (a) An intronic start site in RAI1: a strong start in both cell lines, a burst of nascent RNA that fades within 2–3 kb, and no stable RNA above the host’s background. (b) A cluster of intronic start sites in ZMIZ1: nascent RNA reaches about 5 kb, still without stable RNA. (c) MCTS2, an imprinted retrocopy of MCTS1 nested in HM13: a strong start in both cell lines and nascent RNA running several kilobases into the host’s intron.

### Many annotated intronic long non-coding RNAs lack a start site of their own

These findings bear on the long non-coding RNAs that catalogues place inside introns. Of the 2,344 such genes in GENCODE that lie on the host’s strand outside readthrough genes, 8.9% had a confirmed capped start at their annotated 5′ end in K562, compared with 12.1% of long non-coding RNAs between genes. Both shares are low because most long non-coding RNAs are expressed in few cell types; what matters is the comparison between classes. The difference was clearest among RNAs that are demonstrably expressed, those with RNA-seq reads joining exactly the two ends of one of their own annotated introns, evidence that the RNA is spliced as annotated: in K562, 31% of these intronic RNAs had a confirmed start, against 58% of spliced long non-coding RNAs between genes (Methods). This suggests that, in these two cell lines, many intronic RNAs that are present are made by host transcription, spliced or processed from the host’s precursor RNA, rather than from a promoter of their own; some may have start sites too weak to detect, and the balance may differ in other cell types.

The nested-gene analysis used for validation (Fig. 1c) provides a control for this conclusion. We applied it to long non-coding genes nested in introns of protein-coding genes; those on the opposite strand cannot be pieces of the host’s transcript and so serve as a reference (Additional file 1: Fig. S7). Nested genes of every kind had a start site of their own less often than stand-alone genes of the same type and expression, even protein-coding genes on the host’s strand, whose status as genes is not in doubt; part of this shortfall therefore reflects how expression is measured for genes inside other genes. Long non-coding genes on the host’s strand, however, fell furthest below expectation, below both protein-coding genes on the same strand and long non-coding genes on the opposite strand. Among the few clearly expressed genes, long non-coding genes on the opposite strand had a start site about as often as stand-alone long non-coding genes of the same expression, whereas those on the host’s strand had one about half as often (Additional file 1: Fig. S7). The shortfall of start sites is therefore largest for RNAs that share the host’s strand, as expected if many of them derive from host transcription.

The pair formed by HM13 and MCTS2 illustrates the reverse relationship, in which a nested gene may affect its host (Fig. 6c). MCTS2 is a retrocopy of MCTS1 (a reverse-transcribed copy of its mRNA) that lies in the fourth intron of HM13 and is imprinted, being expressed from the paternal allele [31, 32]. At the corresponding mouse locus, transcription of the nested Mcts2 gene terminates H13 transcription inside the intron [17, 18]. In both human cell lines, MCTS2 had a strong capped start, and nascent RNA from it ran several kilobases into the HM13 intron. A polyadenylation site listed in the PolyASite atlas lies on the HM13 strand about 1.9 kb before the MCTS2 start, and RNA-seq coverage of the intron fell after it, by about 40% in K562 and by half in HCT116. This fits early termination of part of the HM13 transcripts, but coverage also tends to fall along long introns, and our data cannot measure the use of this polyadenylation site directly. The mouse mechanism may not apply in human cells, however: in human fetal tissues, HM13 is not imprinted, and its short isoform that ends before MCTS2 is expressed from both alleles [32], so early termination of HM13 there does not depend on which allele expresses MCTS2. That short isoform ends more than 7 kb before the MCTS2 promoter, at a different site from the PolyASite cluster 1.9 kb before it (Fig. 6c shows only the canonical HM13 transcript). Whether MCTS2 transcription contributes to early termination of HM13 in human cells therefore remains open.

### Inside genes, initiation on the gene’s own strand is weaker

Finally, we asked whether initiation itself is weaker inside genes, which the comparisons above could not show. Most start sites come in divergent pairs, with one start on each strand firing away from the other [8]. Because the two arms of a pair share their surroundings, comparing them controls for local chromatin and mappability. For pairs between genes, which have no gene to define a sense strand, an arm chosen at random was on average as strong as the other arm. Inside introns of active genes, however, the arm on the host gene’s strand was weaker than the opposite arm, by about a third in K562 and by a fifth in HCT116 (Additional file 1: Fig. S8).

The deficit was not an effect of signal depth: it remained when each intronic pair was matched with the pair between genes of the closest total reads (33% in K562 and 17% in HCT116). It also did not scale with host transcription. Pairs in silent genes did not differ significantly from pairs in active genes, and the asymmetry was unrelated, or only very weakly related, to the strength of the host promoter. In silent genes, the arm on the host’s strand also tended to be weaker, but with far fewer pairs this was significant only in HCT116, not in K562, nor when the pairs were matched by depth to pairs between genes. H3K36me3, which the enzyme SETD2 deposits in gene bodies during transcription [33], recruits factors that suppress cryptic initiation [14], so a barrier set up by passing polymerases should act mainly where this mark is present; the deficit was somewhat larger with the mark in both cell lines, but not significantly (Additional file 1: Fig. S8, where all statistics of this section are given). The deficit appeared at every strength of the pairs themselves, whereas pairs between genes stayed symmetric at every strength. Nor was it explained by the excess of G and T that introns carry on the coding strand, which arises because mutation and repair differ between the two strands of transcribed DNA [30]. Initiation on the gene’s own strand is thus weaker inside active genes and probably also inside silent ones, and the deficit does not scale with host transcription, which does not support a simple barrier set up in chromatin by passing host polymerases.

## Discussion

By mapping where RNA polymerase II starts transcription rather than which RNAs are annotated, we found that initiation inside human introns is common but rarely productive. More than a thousand intronic sites in K562 and three thousand in HCT116 carry capped starts. The cap test detects nearly all clearly expressed protein-coding genes nested in introns, and fewer than 1% of random intronic positions pass it, so false start sites are unlikely to explain why so few intronic sites produce stable RNA. Most intronic start sites behave like enhancers or weak promoters: they carry active chromatin marks and the motifs of each cell’s key transcription factors, they mostly initiate in both directions (91% of confirmed sites in K562 and 80% in HCT116 lie in bidirectional PRO-cap peaks), and their transcription mostly stops within a few kilobases, leaving short RNA of which at least part is degraded by the nuclear exosome (Fig. 7b). Intragenic enhancers can act as alternative promoters that produce spliced, polyadenylated RNAs [15], but in the two cell lines we studied, few intronic start sites gave rise to stable RNA.

**Figure 7.**
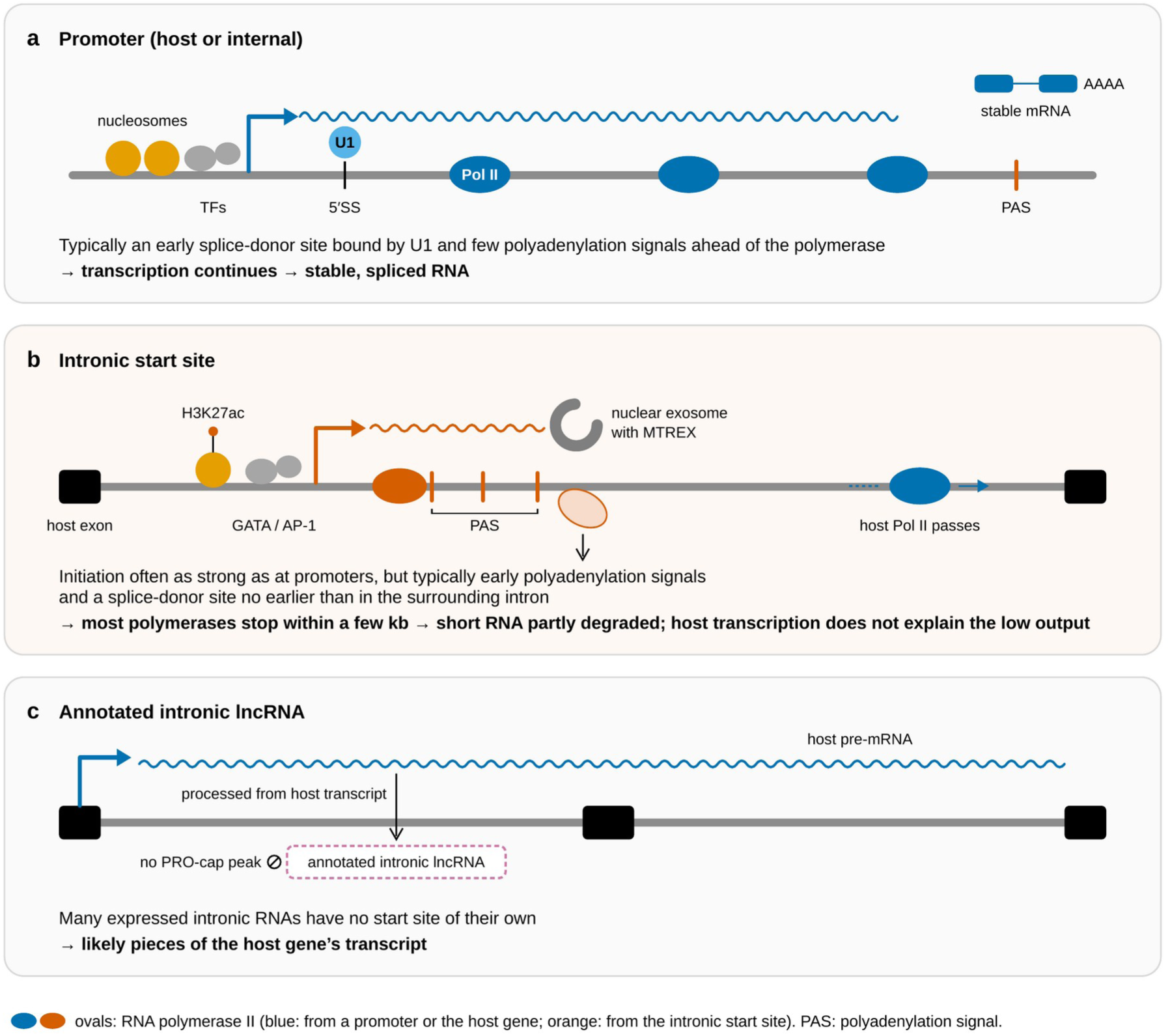
Model: initiation inside introns rarely makes a gene. (a) Promoters (host or internal) typically carry an early splice-donor site bound by U1 and few polyadenylation signals (PAS) ahead of the polymerase, which lets transcription continue and gives stable, spliced RNA. (b) Intronic start sites initiate, often as strongly as promoters, but the sequence downstream resembles the surrounding intron: most polymerases stop within a few kilobases, part of the short RNA is degraded by the nuclear exosome, and sites with more polyadenylation signals respond less after DRB release. Start sites between genes make about as little stable RNA, so host transcription does not explain the low output. (c) Many annotated intronic RNAs that are expressed have no capped start of their own and probably derive from host transcription. Ovals, RNA polymerase II: blue from a promoter or the host gene, orange from the intronic start site.

The low output of intronic start sites is not explained by their location. Per initiation event, intronic start sites in active genes left about as little stable RNA as start sites between genes, which have no host, and their nascent transcription even reached further, although not from sites in silent genes, which lie in the same kind of intronic sequence. The extra reach is therefore more likely due to the transcribed surroundings, such as open chromatin or elongation factors recruited by host transcription, than to the sequence, and it left no more stable RNA. This argues against the idea that host transcription generally suppresses the output of intronic start sites once they have initiated, although interference may still operate at individual loci or at strongly transcribed genes. Inside transcribed genes, cryptic initiation is suppressed by DNA methylation that H3K36me3 recruits in mouse embryonic stem cells [14], and in human cells, depletion of SETD2, the enzyme that deposits H3K36me3, leads to intragenic initiation in at least a tenth of active genes [34]; the sites we map carry the motifs and chromatin marks of active regulatory elements and may escape this control. What separates the host genes’ promoters from intronic start sites is the arrangement downstream of the start: promoters carry early splice-donor sites and few polyadenylation signals in the direction of transcription, whereas intronic start sites resemble the surrounding intronic sequence. This is consistent with the code that governs the choice between productive and PROMPT transcription at promoters [5, 10], here seen at start sites inside genes. The arrangement separates promoters from intronic start sites as groups. At single sites, it did not predict stable RNA, but sites with more polyadenylation signals more often failed to respond after DRB release; the fate of single sites presumably also depends on chromatin, bound factors and the elongation machinery.

The mild shortage of polyadenylation signals on the strand of introns transcribed with the host suggests how new genes could arise inside introns. A regulatory element that already initiates transcription would need to acquire an early splice-donor site and lose nearby polyadenylation signals, as proposed for the birth of genes from divergent transcription at promoters and enhancers [35]; comparisons across mammals show that enhancers have indeed often been converted into promoters during evolution, which likely contributed to the origin of new genes and exons [36]. The small group of start sites shared by both cell lines, which is more conserved and less often derived from transposable elements, is a natural place to look for such cases.

Splicing can also act on initiation. Splicing of internal exons activates weak promoters upstream of them [37, 38], and U1, by suppressing premature polyadenylation, speeds elongation through introns [39] and supports the use of promoters further downstream in the same gene [40]. If splicing of a downstream exon supported initiation, start sites closer to the next exon should initiate more strongly. Because our sites lie at least 1 kb from exons by design, we could test this only from 1 kb upstream of an exon; in this range, the distance to the next exon of the host gene was at most weakly related to how strongly intronic start sites initiated and not related to how often their transcription responded after DRB release (Additional file 1: Fig. S9b,c). Two weaker observations pointed in the direction expected if splicing of the exon supports initiation: initiation tended to be stronger upstream of exons with stronger splice-donor sites in HCT116, though not in K562, and sites upstream of alternatively spliced exons initiated more weakly than sites upstream of constitutive exons in K562, with the same trend in HCT116 (Methods). The support that host transcription lends to downstream promoters [40] may also explain why transcription from intronic sites reached further in active genes.

Our results bear directly on the many long non-coding RNAs that catalogues place inside introns. Expressed intronic RNAs on the host’s strand lack a start site of their own more often than intronic RNAs on the opposite strand or long non-coding RNAs between genes, so a substantial fraction of them are probably processed pieces of the host’s transcript. Some of them may still be genes, but functional studies of such RNAs should first establish an independent start site; otherwise, knockdown or knockout effects may reflect changes to the host gene. Clearly expressed intronic RNAs on the opposite strand had their own start sites about as often as stand-alone long non-coding genes of the same expression, as expected for independent genes.

The HM13/MCTS2 pair points to the other direction of influence between nested and host genes. At the mouse locus, transcription of the nested gene makes transcription of its host end early [18], and transcription at intragenic enhancers can attenuate host transcription [16]. In both human cell lines, MCTS2 also initiates strongly and the host’s RNA coverage falls after a polyadenylation site upstream of it, as the mouse mechanism would predict. In human fetal tissues, however, the short HM13 isoform is made from both alleles, whereas MCTS2 is expressed from one [32], so early termination of HM13 there does not require MCTS2 transcription. Whether MCTS2 contributes to it in human cells could be tested by silencing MCTS2 or by allele-specific 3′-end sequencing. Our genome-wide data leave open whether intronic transcription affects hosts; they show only that hosts in general do not prevent intronic initiation and do not explain its low output.

Inside genes, initiation on the gene’s own strand was weaker than on the opposite strand. This strand asymmetry, measured within divergent pairs of start sites, did not scale with host transcription, did not differ significantly with H3K36me3 and was not explained by base composition, which does not support a simple chromatin barrier, that is, chromatin left behind by passing host polymerases that would block initiation in proportion to how often the gene is transcribed. It may instead reflect a property of the elements themselves, for example features of their sequence that favour initiation on the opposite strand. Whether intronic sequence has also been shaped by selection against sense start sites remains to be tested. If the asymmetry came from the sequence, it should appear in silent genes too. It pointed the same way there, but with few pairs it was significant in only one of the two cell lines, so we report it as an observation rather than as a mechanism.

Our study has limitations. We analysed two cancer cell lines, and HCT116 PRO-cap came from untreated cells of degron lines, without an uncapped control, so the cap test was less strict there; in degron lines, the tagged proteins can also be partly depleted even without treatment. PRO-cap depth also differed (19.1 and 46 million reads), so part of the cell-type specificity of weak sites may reflect detection. We did not correct for multiple testing across the many comparisons; the main conclusions rest on effects with very small p values or on patterns seen in both cell lines, and several rest on the absence of significant differences, which does not exclude small effects. Weak start sites may be missed at the available depth. Start sites inside young transposable elements, whose copies are nearly identical, are also hard to detect, because their reads cannot be mapped uniquely; since random positions were not restricted to uniquely mappable sequence, part of the depletion of transposable elements at start sites may be technical. The MTREX knockout was partial and the RNA-seq libraries captured polyadenylated RNA only, so the contribution of the exosome is underestimated. The DRB data from K562 were not sensitive enough even for the group test (Additional file 1: Fig. S4), and the single-site classification in HCT116 is conservative in strongly transcribed hosts, where the high 40-minute reference signal can hide a burst from the site.

Whether HM13 transcripts are polyadenylated inside the intron could not be tested directly: this needs reads that span the end of the RNA and the start of its poly(A) tail, and the total RNA-seq alignments contained none, presumably because unaligned read ends had been trimmed (Methods). Long-read sequencing of nascent RNA, 3′-end sequencing and targeted perturbation of intronic start sites would test our conclusions at single loci.

## Conclusions

Transcription starts at thousands of sites inside the introns of human protein-coding genes, most of them unannotated and specific to the cell type. Many of these sites initiate as strongly as the hosts’ alternative promoters, yet few give rise to stable RNA: their transcription mostly stops within a few kilobases, their short products are partly degraded by the nuclear exosome, and the sequence downstream rarely has the early splice-donor site that protects promoter transcripts. Start sites between genes leave about as little stable RNA per initiation event, so host transcription does not explain this low output. Expressed intronic RNAs on the host’s strand lack a start site of their own more often than those on the opposite strand, so many of them are probably processed pieces of host transcripts rather than independent genes. Before such RNAs are studied as genes, an independent start site should be established; the intronic start sites listed in Additional file 2 allow a first check in K562 and HCT116 cells.

## Methods

### Data, annotation and processing of nascent RNA

All analyses used the GRCh38 primary assembly and the GENCODE release 47 annotation [2]. Public data came from ENCODE [41, 42] and GEO (accessions in Additional file 1: Table S2). For K562, we used PRO-cap from capped RNA (ENCSR261KBX) and from uncapped RNA as background (ENCSR114HGS); the strand-specific signal files of the two replicates of each were summed with a script that keeps negative values on the minus strand (bigWigMerge discards them). Transcription start sites were taken from the ENCODE peak calls made with PINTS (Peak Identifier for Nascent Transcript Starts) [19], unidirectional (ENCFF636REV) and bidirectional (ENCFF819CPA). For HCT116, we used PRO-cap and PINTS peaks from untreated cells of two degron lines, the CTCF line (ENCSR165QKR) for the main analyses and the RAD21 line (ENCSR635SXX) to check reproducibility; no uncapped control is available for HCT116. Total RNA-seq came from ENCODE (K562 ENCSR792OIJ, HCT116 ENCSR698RPL), as did ChIP-seq peaks for H3K4me3, H3K27ac, H3K4me1 and H3K36me3 in both cell lines, and RNA-seq from K562 cells with CRISPR knockouts of MTREX, EXOSC10 and EXOSC5 and their controls. Polyadenylation sites came from PolyASite 2.0 [43], conservation from the UCSC phyloP track for 100 vertebrates [44], transposable elements from the UCSC RepeatMasker table (RepeatMasker Open-4.0, http://www.repeatmasker.org), and tissue expression from the GTEx v8 table of median expression per tissue [45].

Nascent RNA after DRB release was taken from two studies. For K562, we used BruDRB-seq 10 minutes after DRB washout and Bru-seq without DRB as reference [25] (GSE55534, two replicates each). For HCT116, we used DRB/TTchem-seq from wild-type cells 10, 20, 30 and 40 minutes after release [27] (method: [26]; GSE148844, two replicates). Reads were trimmed with fastp [46], aligned with HISAT2 [47] to an index built from the GENCODE genome, filtered for mapping quality of at least 10 and converted to strand-specific coverage files. Both libraries were sequenced from the strand opposite to the RNA; the scripts detected this automatically from the signal over active host promoters and swapped the strands.

### Intronic start sites, the cap test and controls

Host genes were protein-coding genes. Genes with any transcript tagged as readthrough in GENCODE were excluded as hosts, because a readthrough transcript joins two neighbouring genes and turns the promoter of the downstream gene into an apparent intronic start. Host exons were the union of exons of the host’s coding, nonsense-mediated decay and non-stop decay transcripts; retained-intron transcripts were excluded because they cover whole introns. An intronic start site was the summit of a PRO-cap peak on the host’s strand inside an intron of the host’s reference transcript (MANE Select, otherwise the Ensembl canonical transcript, otherwise the longest) and at least 1 kb from any host exon; summits within 50 nt on the same strand were merged. Peaks were classed as unidirectional, bidirectional or both. Start sites within 500 nt of an annotated intronic long non-coding RNA were linked to it, and start sites within 500 nt of another protein-coding gene’s annotated start were flagged.

A site passed the cap test when it carried at least five capped reads within 100 nt, when this count was significantly higher than expected from the flanks 500–2,000 nt away (Poisson test, p < 0.001, with a background floor of 0.001 reads per nucleotide) and, in K562, when capped reads exceeded uncapped reads at least twofold after normalisation to library depth. The host’s start in use was the annotated start with the most capped reads within 100 nt, and a host gene was considered active when this start carried at least five capped reads. As positive controls, we used the host genes’ internal promoters: annotated alternative starts inside host introns whose transcripts join downstream host exons, at least 1 kb from upstream exons. The two groups barely overlapped: one K562 and two HCT116 intronic start sites lay within 100 nt of an internal promoter. As negative controls, we drew three random positions per site in introns of the same host genes, on the host’s strand and at least 1 kb from exons. Robustness was checked with a 500-nt distance rule instead of 1 kb: in K562 it gave 1,359 instead of 1,100 confirmed sites in active hosts, with a similar chromatin profile (90% with H3K27ac and 53% with H3K4me3, against 89% and 48% in the main analysis) and the same weak response to DRB (8.6% above the control threshold); other window sizes in the DRB test (6 and 4 kb or 10 and 6 kb after and before the site) gave the same result. Reproducibility in HCT116 was assessed by applying the same cap test, at the sites confirmed in the CTCF line, to PRO-cap from untreated cells of the RAD21 degron line.

To validate the approach on known genes, we collected GENCODE protein-coding genes whose whole span lies inside an intron of another protein-coding gene, on either strand and in any of the host’s transcripts, with all annotated starts at least 500 nt from host exons (404 genes: 103 on the host’s strand and 301 on the opposite strand). Stand-alone protein-coding genes not overlapping any other protein-coding gene served as reference. Expression in each cell line was the mean RNA-seq coverage over the gene’s exons on its strand minus the mean coverage of the surrounding intron 2–5 kb on both sides, so that the host’s precursor RNA was not counted. A gene was detected when the cap test passed within 300 nt of any of its annotated starts. Genes were grouped by expression into a bin without signal and fifths of expressed stand-alone genes; the expected number of detected nested genes was the sum of detection rates of stand-alone genes in the same bin. A gene was clearly expressed when its exon signal was at least three times that of its surroundings and in the top two fifths. Well-established genes had a MANE Select transcript [48], and broadly expressed genes had a median of at least 1 TPM in at least 80% of GTEx tissues. Long non-coding genes nested in introns of protein-coding genes were analysed in the same way, separately for those on the host’s strand and on the opposite strand, with stand-alone long non-coding genes as reference (Additional file 1: Fig. S7).

### Chromatin, conservation, transposons and motifs

A site carried a histone mark when an ENCODE peak of that mark lay within 1 kb. Sites with H3K4me3 were called promoter-like, sites with H3K27ac or H3K4me1 but no H3K4me3 enhancer-like, and the rest unmarked. Conservation was the mean phyloP score over 201 nt centred on the site, computed with bigWigAverageOverBed. Transposable elements were taken from the RepeatMasker element covering the start nucleotide, with its class, family and orientation relative to transcription from the site; enrichment was the share of intronic start sites in each class divided by the share of random intronic positions. Transcription factor motifs were tested with HOMER [49] (findMotifsGenome.pl, known motifs only) on 300 nt centred on each intronic start site against the same windows around random intronic positions in the same genes, separately for each cell line.

Start sites of the two cell lines were considered shared when sites on the same strand lay within 200 nt of each other. Shared and cell-type-specific sites with a confirmed start in an active host were compared within thirds of PRO-cap strength, with Fisher’s exact test per third and a Cochran– Mantel–Haenszel test across thirds, which combines the three 2 × 2 tables into a common odds ratio and so compares the groups at equal initiation strength.

### Stable RNA, splicing and the exosome knockouts

Stable RNA after a site was measured in total RNA-seq on the site’s strand as log2 of the mean coverage 0–3 kb after the site divided by the mean coverage 0.2–3.2 kb before it. A site had stable RNA when this value exceeded the 95th percentile of random intronic positions in active hosts. Because the window after a site contains the next exon of the host gene when that exon lies closer than 3 kb, host mRNA can mimic a rise in stable RNA: random intronic positions 1–2 kb upstream of an exon passed this threshold in 26–28% of cases, against at most 0.9% beyond 3 kb (Additional file 1: Fig. S9a). In active hosts, stable RNA was therefore measured only for sites at least 3 kb upstream of the next exon of the host’s reference transcript (84–85% of intronic start sites, 76–78% of internal promoters and 80% of random positions), with the threshold taken from random positions that met the same rule (log2 ratio 0.75 in K562 and 0.99 in HCT116). For the comparison at equal initiation strength, intronic start sites and internal promoters were divided into thirds of PRO-cap reads at the site, with the same boundaries for both groups. For annotated intronic RNAs, splicing was confirmed when at least two RNA-seq reads joined exactly the ends of one of the RNA’s own introns, which the unspliced host precursor RNA cannot produce; junction reads were extracted from the ENCODE alignments. Annotated starts of intronic long non-coding RNAs were tested without the 1-kb distance rule; with it, 3.5% (K562) and 5.1% (HCT116) had a confirmed start. In K562, 72 intronic long non-coding RNAs (5% of those with more than one exon) had splicing evidence, against 7–8% of other classes of long non-coding RNA.

For each site in an active host we measured the distance to the next exon of the host’s reference transcript in the direction of transcription. The splice-donor site of that exon was scored with a position weight matrix built from the donor sites (three exonic and six intronic positions) of all internal exons of the reference transcripts. For sites less than 10 kb upstream of an exon that is not the last, initiation strength was compared across thirds of donor scores (Kruskal–Wallis test; median capped reads from the weakest to the strongest third 43, 46 and 46 in K562, p = 0.9, and 44, 47 and 57 in HCT116, p = 0.07). Inclusion of the exon was estimated from RNA-seq splice junctions (ENCODE junction files in K562, junctions extracted from the ENCSR698RPL alignments in HCT116) as the mean count of the two junctions flanking the exon divided by that mean plus the count of junctions skipping it, for exons with at least 10 junction reads; exons with inclusion below 0.9 were counted as alternative. Sites upstream of alternative exons had a median of 26 capped reads against 49 upstream of constitutive exons in K562 (n = 20 and 315, p = 0.008, Mann–Whitney test) and 32 against 51 in HCT116 (n = 57 and 669, p = 0.11).

For the exosome knockouts, we downloaded for each replicate and strand the ENCODE signal file marked as default (otherwise the newest) and summed the replicates. Each knockout was first checked with a positive control: the rise of upstream antisense RNAs (PROMPTs), measured as log2(knockout/control) 200–2,000 nt upstream of active host promoters on the opposite strand, minus the same ratio over the first 3 kb of the gene. The effect at each site was the change in the stable RNA measure above between knockout and control. Sites without any reads in the four windows of the knockout and control libraries were excluded, and in active hosts the 3-kb rule above was applied, with the threshold from random positions meeting the same rule; this left 523 of the 1,100 K562 intronic start sites in active hosts. Sites counted as responsive above the 95th percentile of random intronic positions, and groups were compared with random positions by a one-sided Mann–Whitney test. Only the MTREX knockout passed the PROMPT check (Additional file 1: Fig. S5); the libraries contain polyadenylated RNA only.

### Nascent transcription after DRB release

The group test compared, for each site, the rise in nascent RNA after the site with the rise in the reference sample: log2[(test after + 0.5)/(test before + 0.5)] − log2[(reference after + 0.5)/(reference before + 0.5)], computed from read counts 0–8 kb after the site and 0.2–5.2 kb before it. The reference was Bru-seq without DRB in K562 and the 40-minute sample in HCT116. In HCT116 only sites at least 40 kb (10 minutes) or 70 kb (20 minutes) from the start in use of an active host were tested, and in K562 sites at least 20 kb from it, so that they lay beyond the front of the host’s own wave; the front was read from the mean nascent RNA profile along active host genes: the signal fell from a quarter of its plateau to background 12–20 kb from the gene start after 10 minutes and 44– 53 kb after 20 minutes (Additional file 1: Fig. S3). Of 1,235 intronic start sites in active hosts beyond this distance, 1,191 had enough reads to be tested; 15.2% of them responded (internal promoters: 19.6% of 194). Without the 38 sites that had an active internal promoter up to 20 kb upstream, whose polymerases could reach the site within 10 minutes, the share was 15.6%. Groups were compared with random intronic positions, and the share of responsive sites was the share above their 95th percentile. Host exons inside the windows did not bias this test: random positions responded at 3–6% at every distance to the next host exon in both cell lines (Additional file 1: Fig. S9c).

Average profiles show, for sites beyond the front, the nascent RNA at 10 or 20 minutes summed over all sites of a group and divided by the summed signal at 40 minutes, in 1-kb bins from 10 kb before to 10 kb after the site, on the site’s strand.

Single sites were classified from the two HCT116 replicates separately. For each replicate, a near score and a far score were computed as above with a window 0.2–5.2 kb before the site and windows 0–2 kb (near) and 2–8 kb (far) after it, comparing 10 minutes with 40 minutes. Thresholds were the 95th percentiles of random intronic positions in active hosts, separately for each replicate and window. A site was productive when the far score exceeded the threshold in both replicates, short when only the near score did so in both, unresponsive when neither did, and unresolved when either replicate had fewer than 20 reads in the windows. Sites qualified when their host was silent or when they lay more than 22.2 kb from the host’s start in use, beyond the 10-minute front. The false discovery rate of each class was estimated from the share of random positions in that class.

### Downstream sequence, start sites between genes and RNA per initiation

For each site, we counted polyadenylation signals (AATAAA or ATTAAA, the AWTAAA motif of reference [5]; overlapping matches allowed) and splice-donor motifs (consensus [AC]AG| GT[AG]AGT) in the first kilobase on the transcribed strand and, for comparison, in the first kilobase in the opposite direction from the same site. The ratio of polyadenylation signals on the two strands was computed per group from mean densities, which controls for local base composition. A site was protected when the first splice-donor motif came before the first polyadenylation signal, or when there was no signal.

Start sites between genes were PRO-cap peak summits at least 5 kb from any protein-coding gene on either strand that passed the same cap test. For the comparison with intronic start sites, all groups were restricted to unannotated sites, more than 500 nt from any annotated transcript start, and divided into thirds of PRO-cap reads defined on the intronic and intergenic groups together. RNA per initiation event was the mean RNA-seq coverage 0–3 kb after the site minus the mean coverage 0.2–3.2 kb before it, divided by the PRO-cap reads at the site (reported ×10⁶); in active hosts, intronic start sites and internal promoters followed the 3-kb rule above. Nascent RNA per initiation was the excess of 10-minute signal 2–8 kb after the site over the level before it, divided in the same way, for sites more than 22.2 kb from the host’s start in use, as for single sites; with the 40-kb rule of the group test, the median for intronic start sites in active hosts was almost unchanged (0.75 against 0.72 × 10⁻³; n = 1,202). We report the median with a 95% bootstrap confidence interval (1,000 resamples) and the share of sites with a positive excess with a Wilson confidence interval.

Divergent pairs were taken from the bidirectional PINTS peaks, with one arm on each strand; in K562, these peaks were 111–265 nt wide (interquartile range). Pairs inside a host intron with both arms at least 1 kb from exons were assigned the host’s strand as sense, and pairs at least 5 kb from protein-coding genes were given an arbitrary orientation as reference. The asymmetry of a pair was log2[(capped reads within 50 nt of the sense arm + 1)/(capped reads within 50 nt of the antisense arm + 1)]. We related it to host activity, to the strength of the host’s promoter, to an H3K36me3 peak within 1 kb and to strand differences in base composition. To exclude an effect of signal depth, each intronic pair was matched to the pair between genes with the closest summed reads of both arms (greedy nearest-neighbour matching without replacement, intronic pairs in random order, ties resolved towards the deeper pair, at most 10% apart; all intronic pairs matched, 93–96% of them with identical depth). The deficit of the sense arm was 1 − 2^d, where d is the difference between the median asymmetries of the intronic pairs and their matched pairs. It was tested with the Wilcoxon signed-rank test on the differences within matched pairs, and its 95% confidence interval comes from 10,000 bootstrap resamples of whole host genes, each intronic pair resampled together with its matched pair, because pairs in the same gene are not independent. Many pairs between genes have the same depth, so the estimate depends somewhat on which of them is chosen: across 200 random matchings within the same 10% window, the middle 95% of the estimates lay between 31% and 43% in K562 and between 15% and 26% in HCT116.

### HM13 and MCTS2, locus figures and statistics

To look for early polyadenylation of HM13, we searched the total RNA-seq alignments for reads ending in at least eight soft-clipped nucleotides of which at least 85% were A, discarding ends followed in the genome by an A-rich stretch (at least seven A in 10 nt), which indicates internal priming. No such reads were found in either cell line, presumably because the alignments do not keep unaligned read ends. We therefore report the PolyASite clusters and the annotated 3′ ends of HM13 transcripts, and RNA-seq coverage on the HM13 strand in 500-nt bins across the intron that holds MCTS2.

Locus figures were drawn with pyGenomeTracks 3.9 [50] in windows from 10 kb before to 20 kb after each site in the direction of transcription (nested genes: the gene ± 10 kb). For display only, PRO-cap reads were summed in 301-nt sliding windows so that narrow start peaks remain visible at this scale. Intronic start sites and internal promoters were chosen among sites with a confirmed start in both cell lines and an active host in both, ranked by the smaller of their two PRO-cap counts.

Groups of sites were compared with Mann–Whitney tests, proportions with Fisher’s exact test and stratified proportions with the Cochran–Mantel–Haenszel test. The asymmetry of divergent pairs was tested against zero with the Wilcoxon signed-rank test, and correlations are Spearman coefficients. Confidence intervals of medians come from 1,000 bootstrap resamples and those of Mantel–Haenszel odds ratios from the Robins–Breslow–Greenland variance. Wherever a single threshold was needed, it was set empirically at the 95th percentile of random intronic positions in the same analysis, so that about 5% of random positions pass by construction. Analyses were written in Python 3.11.16 with pandas 3.0.6, NumPy 2.4.6, SciPy 1.17.1 and pyBigWig 0.3.26 (https://doi.org/10.5281/zenodo.594045), together with samtools 1.24 [51], bedtools 2.31.1 [52], fastp 1.3.7, HISAT2 2.2.3, HOMER 5.1 and the UCSC tools bigWigAverageOverBed and bigWigMerge, release 482 [53]. Comparisons of groups with random intronic positions in the DRB and exosome analyses were one-sided, testing for a greater response than at random positions, because only a response above that of random positions was of interest; all other tests were two-sided. P values were not adjusted for multiple comparisons; the conclusions rest on effects with very small p values or on patterns seen in both cell lines.

### Use of generative AI tools

Claude (Anthropic) was used to help write and debug analysis code, cross-check numbers in the text against the analysis output, and edit the text. The authors ran and verified all code, checked all text, numbers and citations, and take full responsibility for the content of the article.

## Abbreviations

BruDRB-seq: bromouridine sequencing after release from DRB
Bru-seq: bromouridine sequencing
ChIP-seq: chromatin immunoprecipitation followed by sequencing
CRISPR: clustered regularly interspaced short palindromic repeats
DRB: 5,6-dichloro-1-β-D-ribofuranosylbenzimidazole
ENCODE: Encyclopedia of DNA Elements
GEO: Gene Expression Omnibus
GTEx: Genotype-Tissue Expression
LTR: long terminal repeat
MANE: Matched Annotation from NCBI and EMBL-EBI
NEXT: nuclear exosome targeting complex
PAS: polyadenylation signal
PINTS: Peak Identifier for Nascent Transcript Starts
PRO-cap: precision nuclear run-on and sequencing of capped RNA
PROMPT: promoter upstream transcript
PSNC: Poznan Supercomputing and Networking Center
RNA-seq: RNA sequencing
TPM: transcripts per million
TTchem-seq: transient transcriptome sequencing with chemical RNA fragmentation
U1: U1 small nuclear ribonucleoprotein
UCSC: University of California, Santa Cruz

## Declarations

### Ethics approval and consent to participate

Not applicable.

### Consent for publication

Not applicable.

### Availability of data and materials

All data analysed in this study are public; accession numbers are listed in Additional file 1: Table S2. Additional file 2: Table S1 lists all 5,801 intronic start sites of both cell lines with every measure used in the paper, including the distance to the next exon of the host gene; stable RNA measures are given only for sites at least 3 kb upstream of that exon. The analysis code (steps 01–45, the batch jobs and the figure scripts) is available at https://github.com/michal-szczesniak/intronic-start-sites and, together with the processed tables, in Zenodo (https://doi.org/10.5281/zenodo.23984488).

Project name: intronic-start-sites. Project home page: https://github.com/michal-szczesniak/intronic-start-sites. Archived version: https://doi.org/10.5281/zenodo.23984488. Operating system: Linux. Programming languages: Python and Bash. Other requirements: Python 3, HOMER, samtools, bedtools, HISAT2, fastp and the UCSC tools (versions in the Methods; conda environments in the repository). License: MIT (code); CC BY 4.0 (processed data). Any restrictions to use by non-academics: none.

### Authors’ contributions

MWS conceived the study, designed the methodology, wrote the software, performed the formal analysis and investigation, curated the data, prepared the visualizations, wrote the original draft, reviewed and edited the manuscript, supervised the work and administered the project. EW conceived the study, wrote the original draft, reviewed and edited the manuscript and acquired funding. Both authors read and approved the final manuscript.

## Acknowledgements

We thank the ENCODE Consortium and the laboratories that produced the public data analysed here. Computations were performed at the Poznan Supercomputing and Networking Center (PSNC).

## Supplementary information

**Additional file 1** (PDF). Figures S1–S10 and Table S2. The supplementary figures with their legends (S1, calibration of the cap test; S2, reproducibility between two HCT116 lines; S3 and S4, nascent RNA after DRB release; S5, exosome knockouts; S6, host precursor RNA and the rise in stable RNA; S7, start sites of nested genes by strand; S8, strand asymmetry of divergent start pairs; S9, distance to the next exon; S10, reference loci), the description of the columns of Table S1, and Table S2, the data sources with their accession numbers.

**Additional file 2** (XLSX). Table S1. All intronic start sites of both cell lines, one row per site and cell line, with every measure used in the paper; a second sheet describes the columns.

## References

1. Derrien T, Johnson R, Bussotti G, et al. The GENCODE v7 catalog of human long noncoding RNAs: analysis of their gene structure, evolution, and expression. Genome Res. 2012;22:1775–89. 10.1101/gr.132159.111

2. Frankish A, Carbonell-Sala S, Diekhans M, et al. GENCODE: reference annotation for the human and mouse genomes in 2023. Nucleic Acids Res. 2023;51:D942–9. 10.1093/nar/gkac1071

3. Kumar A. An overview of nested genes in eukaryotic genomes. Eukaryot Cell. 2009;8:1321–9. 10.1128/EC.00143-09

4. Montibus B, Cain JA, Martinez-Nunez RT, et al. Global identification of mammalian host and nested gene pairs reveal tissue-specific transcriptional interplay. Genome Res. 2024;34:2163–75. 10.1101/gr.279430.124

5. Ntini E, Järvelin AI, Bornholdt J, et al. Polyadenylation site–induced decay of upstream transcripts enforces promoter directionality. Nat Struct Mol Biol. 2013;20:923–8. 10.1038/nsmb.2640

6. Preker P, Nielsen J, Kammler S, et al. RNA exosome depletion reveals transcription upstream of active human promoters. Science. 2008;322:1851–4. 10.1126/science.1164096

7. Kim TK, Hemberg M, Gray JM, et al. Widespread transcription at neuronal activity-regulated enhancers. Nature. 2010;465:182–7. 10.1038/nature09033

8. Core LJ, Martins AL, Danko CG, et al. Analysis of nascent RNA identifies a unified architecture of initiation regions at mammalian promoters and enhancers. Nat Genet. 2014;46:1311–20. 10.1038/ng.3142

9. Andersson R, Gebhard C, Miguel-Escalada I, et al. An atlas of active enhancers across human cell types and tissues. Nature. 2014;507:455–61. 10.1038/nature12787

10. Almada AE, Wu X, Kriz AJ, et al. Promoter directionality is controlled by U1 snRNP and polyadenylation signals. Nature. 2013;499:360–3. 10.1038/nature12349

11. Kaida D, Berg MG, Younis I, et al. U1 snRNP protects pre-mRNAs from premature cleavage and polyadenylation. Nature. 2010;468:664–8. 10.1038/nature09479

12. Schlackow M, Nojima T, Gomes T, et al. Distinctive patterns of transcription and RNA processing for human lincRNAs. Mol Cell. 2017;65:25–38. 10.1016/j.molcel.2016.11.029

13. Shearwin KE, Callen BP, Egan JB. Transcriptional interference – a crash course. Trends Genet. 2005;21:339–45. 10.1016/j.tig.2005.04.009

14. Neri F, Rapelli S, Krepelova A, et al. Intragenic DNA methylation prevents spurious transcription initiation. Nature. 2017;543:72–7. 10.1038/nature21373

15. Kowalczyk MS, Hughes JR, Garrick D, et al. Intragenic enhancers act as alternative promoters. Mol Cell. 2012;45:447–58. 10.1016/j.molcel.2011.12.021

16. Cinghu S, Yang P, Kosak JP, et al. Intragenic enhancers attenuate host gene expression. Mol Cell. 2017;68:104–17.e6. 10.1016/j.molcel.2017.09.010

17. Wood AJ, Schulz R, Woodfine K, et al. Regulation of alternative polyadenylation by genomic imprinting. Genes Dev. 2008;22:1141–6. 10.1101/gad.473408

18. Cain JA, Yang C, Kolbeinsdottir H, et al. Transcriptional interference drives intronic polyadenylation at the endogenous H13/Mcts2 locus. Nucleic Acids Res. 2026;54:gkag640. 10.1093/nar/gkag640

19. Yao L, Liang J, Ozer A, et al. A comparison of experimental assays and analytical methods for genome-wide identification of active enhancers. Nat Biotechnol. 2022;40:1056–65. 10.1038/s41587-022-01211-7

20. Kwak H, Fuda NJ, Core LJ, et al. Precise maps of RNA polymerase reveal how promoters direct initiation and pausing. Science. 2013;339:950–3. 10.1126/science.1229386

21. Jang HS, Shah NM, Du AY, et al. Transposable elements drive widespread expression of oncogenes in human cancers. Nat Genet. 2019;51:611–7. 10.1038/s41588-019-0373-3

22. Brocks D, Schmidt CR, Daskalakis M, et al. DNMT and HDAC inhibitors induce cryptic transcription start sites encoded in long terminal repeats. Nat Genet. 2017;49:1052–60. 10.1038/ng.3889

23. Gutiérrez L, Caballero N, Fernández-Calleja L, et al. Regulation of GATA1 levels in erythropoiesis. IUBMB Life. 2020;72:89–105. 10.1002/iub.2192

24. McConnell BB, Kim SS, Yu K, et al. Krüppel-like factor 5 is important for maintenance of crypt architecture and barrier function in mouse intestine. Gastroenterology. 2011;141:1302–13.e6. 10.1053/j.gastro.2011.06.086

25. Veloso A, Kirkconnell KS, Magnuson B, et al. Rate of elongation by RNA polymerase II is associated with specific gene features and epigenetic modifications. Genome Res. 2014;24:896–905. 10.1101/gr.171405.113

26. Gregersen LH, Mitter R, Svejstrup JQ. Using TTchem-seq for profiling nascent transcription and measuring transcript elongation. Nat Protoc. 2020;15:604–27. 10.1038/s41596-019-0262-3

27. Geijer ME, Zhou D, Selvam K, et al. Elongation factor ELOF1 drives transcription-coupled repair and prevents genome instability. Nat Cell Biol. 2021;23:608–19. 10.1038/s41556-021-00692-z

28. Lubas M, Christensen MS, Kristiansen MS, et al. Interaction profiling identifies the human nuclear exosome targeting complex. Mol Cell. 2011;43:624–37. 10.1016/j.molcel.2011.06.028

29. Proudfoot NJ. Ending the message: poly(A) signals then and now. Genes Dev. 2011;25:1770–82. 10.1101/gad.17268411

30. Green P, Ewing B, Miller W, et al. Transcription-associated mutational asymmetry in mammalian evolution. Nat Genet. 2003;33:514–7. 10.1038/ng1103

31. Wood AJ, Roberts RG, Monk D, et al. A screen for retrotransposed imprinted genes reveals an association between X chromosome homology and maternal germ-line methylation. PLoS Genet. 2007;3:e20. 10.1371/journal.pgen.0030020

32. Monk D, Arnaud P, Frost JM, et al. Human imprinted retrogenes exhibit non-canonical imprint chromatin signatures and reside in non-imprinted host genes. Nucleic Acids Res. 2011;39:4577–86. 10.1093/nar/gkq1230

33. Markert JW, Soffers JH, Farnung L. Structural basis of H3K36 trimethylation by SETD2 during chromatin transcription. Science. 2025;387:528–33. 10.1126/science.adn6319

34. Carvalho S, Raposo AC, Martins FB, et al. Histone methyltransferase SETD2 coordinates FACT recruitment with nucleosome dynamics during transcription. Nucleic Acids Res. 2013;41:2881–93. 10.1093/nar/gks1472

35. Wu X, Sharp PA. Divergent transcription: a driving force for new gene origination? Cell. 2013;155:990–6. 10.1016/j.cell.2013.10.048

36. Carelli FN, Liechti A, Halbert J, et al. Repurposing of promoters and enhancers during mammalian evolution. Nat Commun. 2018;9:4066. 10.1038/s41467-018-06544-z

37. Fiszbein A, Krick KS, Begg BE, et al. Exon-mediated activation of transcription starts. Cell. 2019;179:1551–65.e17. 10.1016/j.cell.2019.11.002

38. Uriostegui-Arcos M, Mick ST, Shi Z, et al. Splicing activates transcription from weak promoters upstream of alternative exons. Nat Commun. 2023;14:3435. 10.1038/s41467-023-39200-2

39. Mimoso CA, Adelman K. U1 snRNP increases RNA Pol II elongation rate to enable synthesis of long genes. Mol Cell. 2023;83:1264–79.e10. 10.1016/j.molcel.2023.03.002

40. Kim G, Carroll CL, Wakefield ZP, et al. U1 snRNP regulates alternative promoter activity by inhibiting premature polyadenylation. Mol Cell. 2025;85:1968–81.e7. 10.1016/j.molcel.2025.04.021

41. ENCODE Project Consortium. Expanded encyclopaedias of DNA elements in the human and mouse genomes. Nature. 2020;583:699–710. 10.1038/s41586-020-2493-4

42. Luo Y, Hitz BC, Gabdank I, et al. New developments on the Encyclopedia of DNA Elements (ENCODE) data portal. Nucleic Acids Res. 2020;48:D882–9. 10.1093/nar/gkz1062

43. Herrmann CJ, Schmidt R, Kanitz A, et al. PolyASite 2.0: a consolidated atlas of polyadenylation sites from 3′ end sequencing. Nucleic Acids Res. 2020;48:D174–9. 10.1093/nar/gkz918

44. Pollard KS, Hubisz MJ, Rosenbloom KR, et al. Detection of nonneutral substitution rates on mammalian phylogenies. Genome Res. 2010;20:110–21. 10.1101/gr.097857.109

45. GTEx Consortium. The GTEx Consortium atlas of genetic regulatory effects across human tissues. Science. 2020;369:1318–30. 10.1126/science.aaz1776

46. Chen S, Zhou Y, Chen Y, et al. fastp: an ultra-fast all-in-one FASTQ preprocessor. Bioinformatics. 2018;34:i884–90. 10.1093/bioinformatics/bty560

47. Kim D, Paggi JM, Park C, et al. Graph-based genome alignment and genotyping with HISAT2 and HISAT-genotype. Nat Biotechnol. 2019;37:907–15. 10.1038/s41587-019-0201-4

48. Morales J, Pujar S, Loveland JE, et al. A joint NCBI and EMBL-EBI transcript set for clinical genomics and research. Nature. 2022;604:310–5. 10.1038/s41586-022-04558-8

49. Heinz S, Benner C, Spann N, et al. Simple combinations of lineage-determining transcription factors prime cis-regulatory elements required for macrophage and B cell identities. Mol Cell. 2010;38:576–89. 10.1016/j.molcel.2010.05.004

50. Lopez-Delisle L, Rabbani L, Wolff J, et al. pyGenomeTracks: reproducible plots for multivariate genomic datasets. Bioinformatics. 2021;37:422–3. 10.1093/bioinformatics/btaa692

51. Danecek P, Bonfield JK, Liddle J, et al. Twelve years of SAMtools and BCFtools. GigaScience. 2021;10:giab008. 10.1093/gigascience/giab008

52. Quinlan AR, Hall IM. BEDTools: a flexible suite of utilities for comparing genomic features. Bioinformatics. 2010;26:841–2. 10.1093/bioinformatics/btq033

53. Kent WJ, Zweig AS, Barber G, et al. BigWig and BigBed: enabling browsing of large distributed datasets. Bioinformatics. 2010;26:2204–7. 10.1093/bioinformatics/btq351

