## Supplemental File 1 for "Transcription starting inside human introns is widespread but mostly yields short-lived RNA"

### Additional file 1

This file contains Figures S1–S10 with their legends, the description of Table S1 (the table itself is Additional file 2, an Excel file) and Table S2.

#### Figures

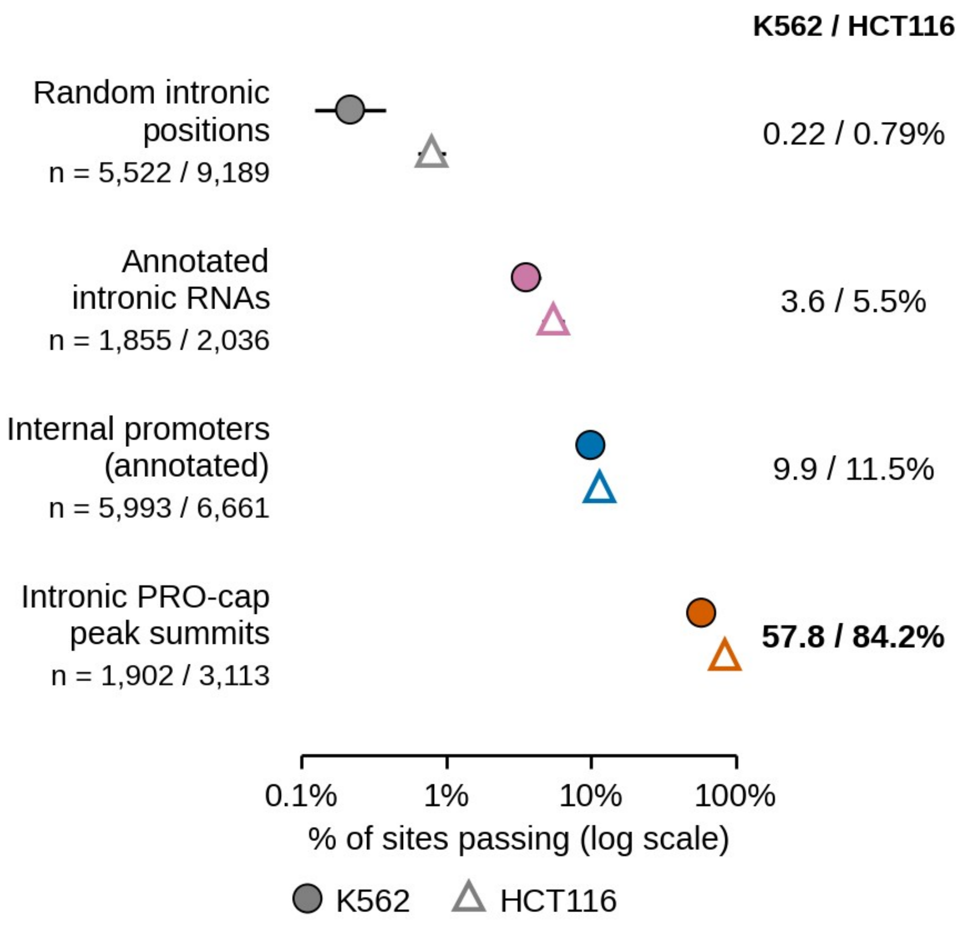


**Figure S1. Calibration of the cap test.** Share of sites passing the test in active host genes, with Wilson 95% confidence intervals, for random positions in the same introns (negative control), annotated intronic RNAs, annotated internal promoters of the host genes and the summits of PRO-cap peaks inside introns. Fewer than 1% of random positions passed. About a tenth of annotated internal promoters passed (595 of 5,993 in K562 and 765 of 6,661 in HCT116): peak summits were selected as PRO-cap peaks in the first place and mostly pass, whereas annotated starts are tested whatever their signal, and each cell type uses only a small fraction of them. Peak summits passed more often in HCT116, probably because no uncapped control exists there, so enrichment over uncapped RNA could not be required. In silent hosts, 0.4% (K562) and 0.3% (HCT116) of random positions passed; no internal promoter can pass there, because a host with a passing annotated start counts as active.

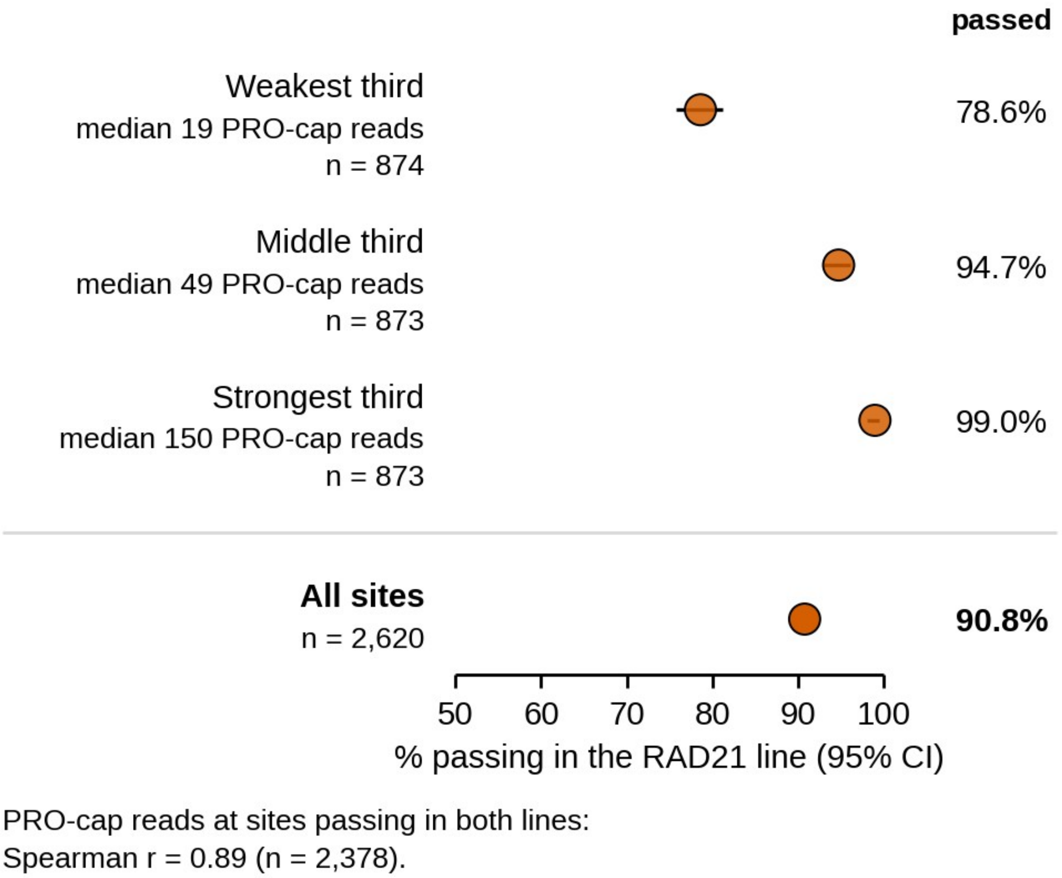


**Figure S2. Intronic start sites are reproducible between two HCT116 lines.** Confirmed intronic start sites in active hosts of the CTCF degron line (main data set) were tested with the same cap test in PRO-cap from untreated cells of the RAD21 degron line, by thirds of PRO-cap strength in the CTCF line. Overall, 91% passed in the RAD21 line, rising from 79% of the weakest third to 99% of the strongest; at sites passing in both, PRO-cap reads correlated closely between the lines (Spearman $\rho = 0.89$, $n = 2{,}378$, $p < 10^{-300}$). Comparing independent peak calls instead, 60% of intronic start sites of the CTCF line were also called as peaks in the RAD21 line.

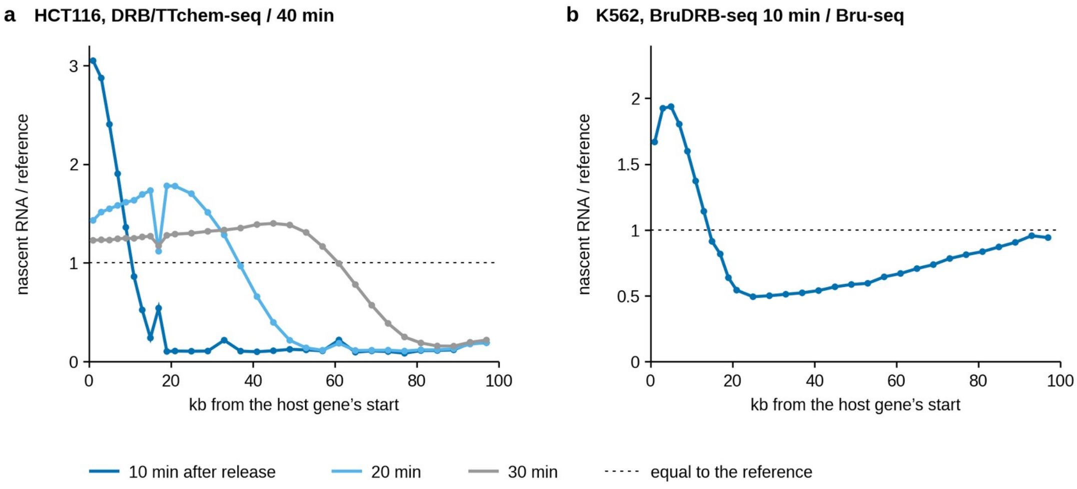


**Figure S3. The wave of transcription after DRB release.** Nascent RNA summed over 1,000 active host genes extending at least 100 kb beyond their start in use, in 2-kb bins (beyond 20 kb, every second bin is shown), divided by the reference sample (40 min after release in HCT116, untreated cells in K562); the dotted line at 1 marks the reference level. (a) HCT116, 10, 20 and 30 min after release relative to 40 min. The front, from where the signal had fallen to a quarter of its plateau to where it reached background, lay 12–20 kb from the gene start after 10 min, 44–53 kb after 20 min and 74–85 kb after 30 min. Beyond it, the signal was about a tenth of the 40-min level: DRB had stopped transcription almost completely. (b) K562, BruDRB-seq 10 min after release relative to Bru-seq. The signal fell below the untreated level at about 14 kb but stayed at about half of it and rose again further into long genes, probably because DRB inhibited transcription incompletely in these data, so the test was much less sensitive (Fig. S4).

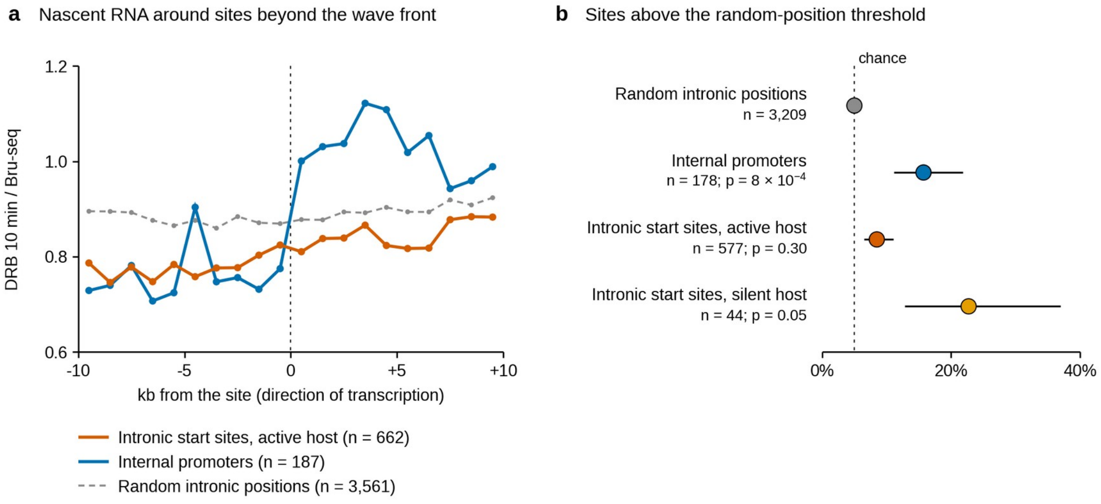


**Figure S4. The DRB test in K562 is not sensitive enough for intronic start sites.** (a) BruDRB-seq 10 min after release divided by Bru-seq, summed over sites beyond the front of the host's wave, in 1-kb bins. Internal promoters show a step after the site; intronic start sites in active hosts rise only slightly, and random positions stay flat. (b) Share of sites with enough reads to be tested (577 intronic start sites and 178 internal promoters in active hosts, 44 intronic start sites in silent hosts and 3,209 random positions) whose rise exceeds the 95th percentile of random positions, with Wilson 95% confidence intervals; p values from one-sided Mann–Whitney tests against random positions. The weak response of intronic start sites (8.5%, against 5% by chance) fits the small burst seen in HCT116, given the high background in K562 (Fig. S3b). Intronic start sites in silent hosts, which have no host background, responded more often.

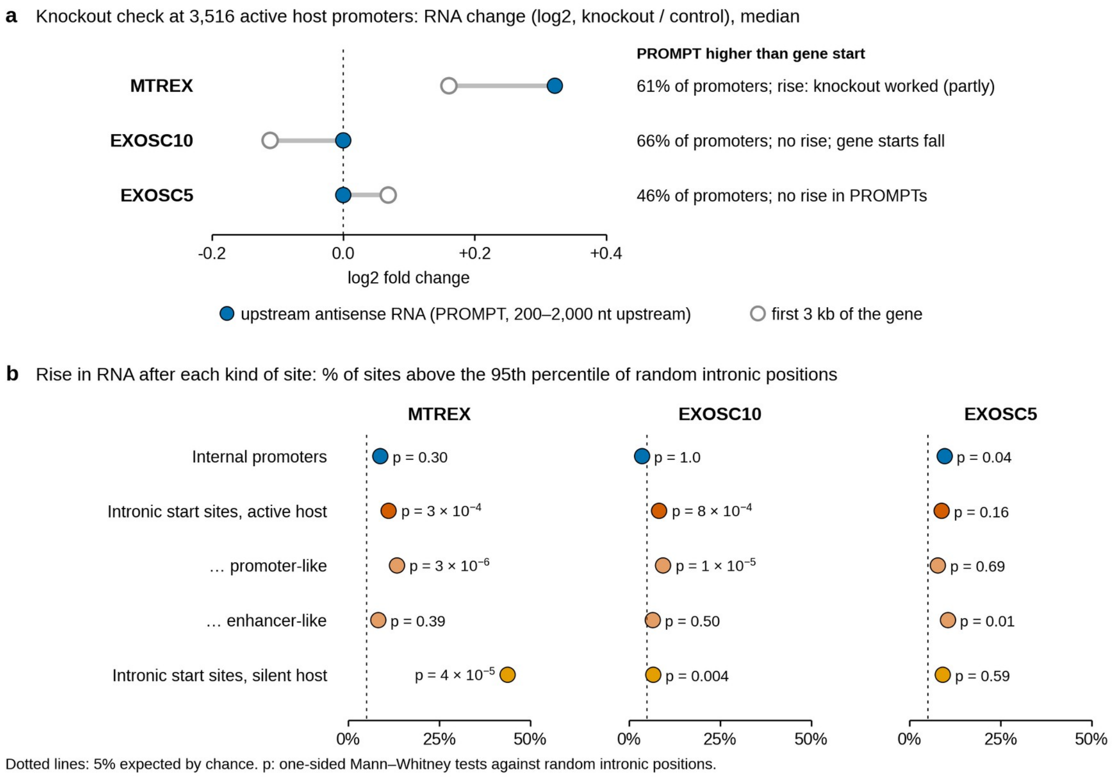


**Figure S5. Only the MTREX knockout raised exosome substrates.** (a) Each ENCODE knockout was checked with upstream antisense RNAs (PROMPTs), classic exosome substrates: median log2 change of RNA 200–2,000 nt upstream of 3,516 active host promoters (opposite strand) and over the first 3 kb of the gene. In MTREX knockout cells, PROMPTs rose (+0.32) more than gene starts (+0.16). In EXOSC10 knockout cells, PROMPTs did not change and the apparent difference came from a fall at gene starts; the EXOSC5 knockout did not change PROMPTs either. (b) Share of sites whose stable RNA rise in knockout relative to control cells exceeded the 95th percentile of random intronic positions, for all sites regardless of the distance to the next host exon (hence MTREX values slightly different from Fig. 4a, which applies the 3-kb rule). Only MTREX is interpreted in the main text; the EXOSC5 knockout, which did not raise PROMPTs, still gave p = 0.01–0.04 for two groups, the level of chance findings here.

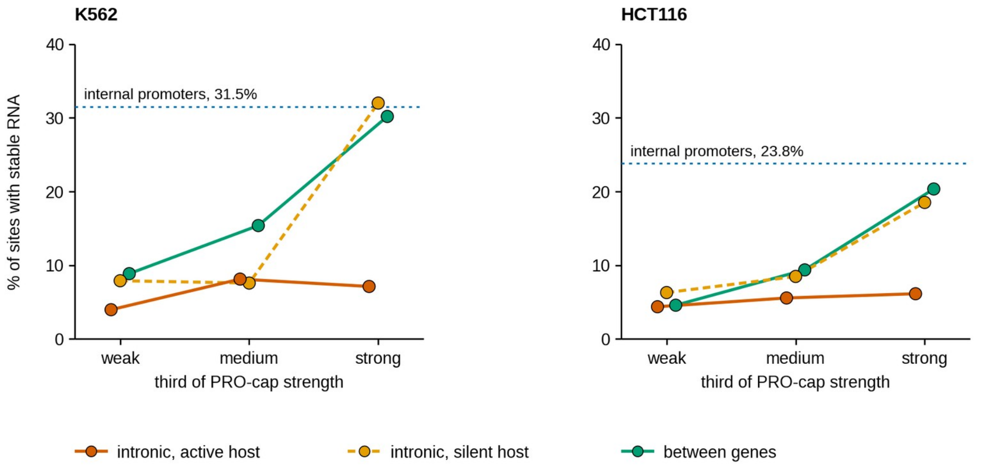


**Figure S6. Host precursor RNA dilutes the rise in stable RNA after intronic sites.** Share of unannotated sites passing the cap test whose stable RNA rises after the site (log2 of coverage 0–3 kb after over 0.2–3.2 kb before) above the 95th percentile of random intronic positions, by thirds of PRO-cap strength, for all sites regardless of the distance to the next host exon (not comparable with Fig. 3a). By this ratio, intronic sites in active hosts look several times weaker than sites between genes or in silent hosts (strongest third: 7% against 30% and 32% in K562; 6% against 20% and 19% in HCT116), because the stretch before the site is already covered by host precursor RNA. Per initiation event, intronic sites in active hosts produce about as little stable RNA as sites between genes (Fig. 5a). Dotted lines: internal promoters, all strengths pooled.

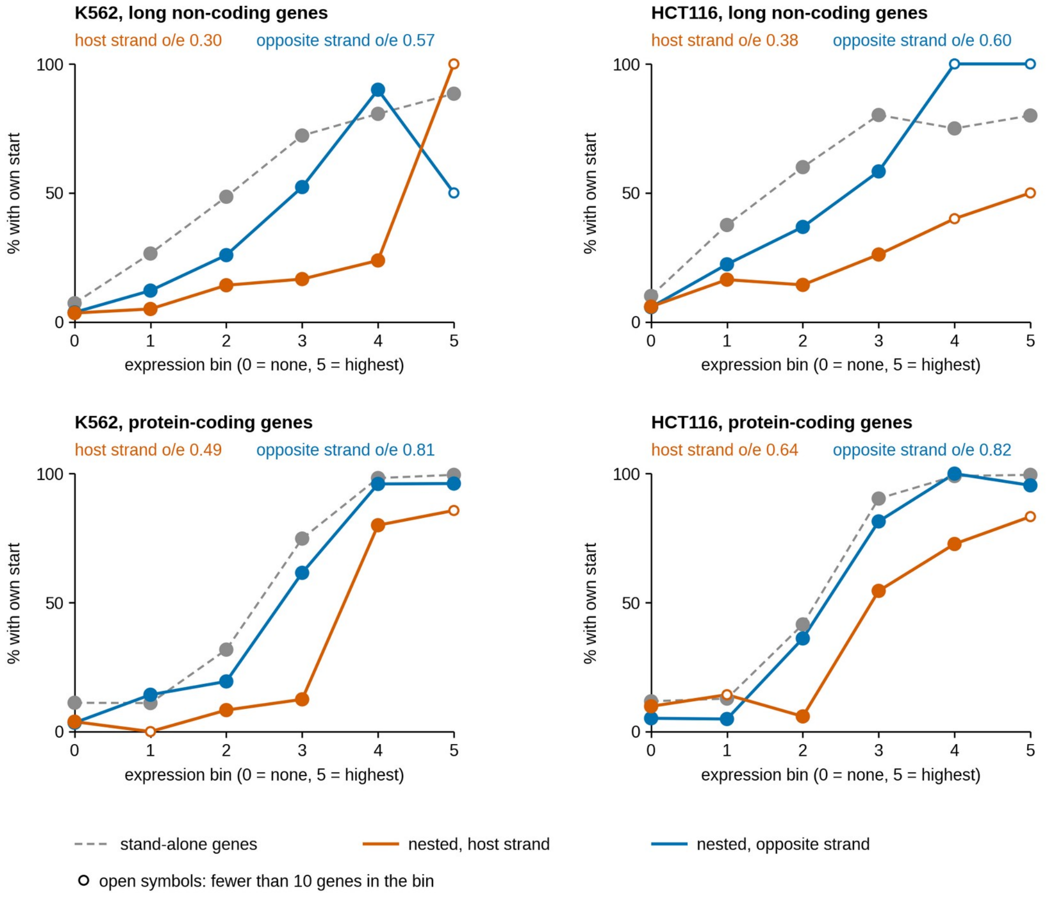

**Figure S7. Start sites of genes nested in introns, by strand.** Share of genes whose annotated start passed the cap test within 300 nt, by expression bin of the gene itself (0, no signal above the surroundings; 1–5, fifths of expressed stand-alone genes), for stand-alone genes and for genes nested in introns of protein-coding genes on the host's strand or on the opposite strand; long non-coding genes (top) and protein-coding genes (bottom), K562 (left) and HCT116 (right). Observed/expected (o/e) over all bins, with the expectation taken from stand-alone genes of the same type and bin, is given in each panel. Open symbols: fewer than 10 genes in the bin. Among clearly expressed long non-coding genes, 18 of 21 (K562) and 7 of 7 (HCT116) on the opposite strand had a start site, against 7 of 14 and 2 of 6 on the host's strand ($p = 0.053$ and 0.021 for the difference between strands, Fisher's exact test).

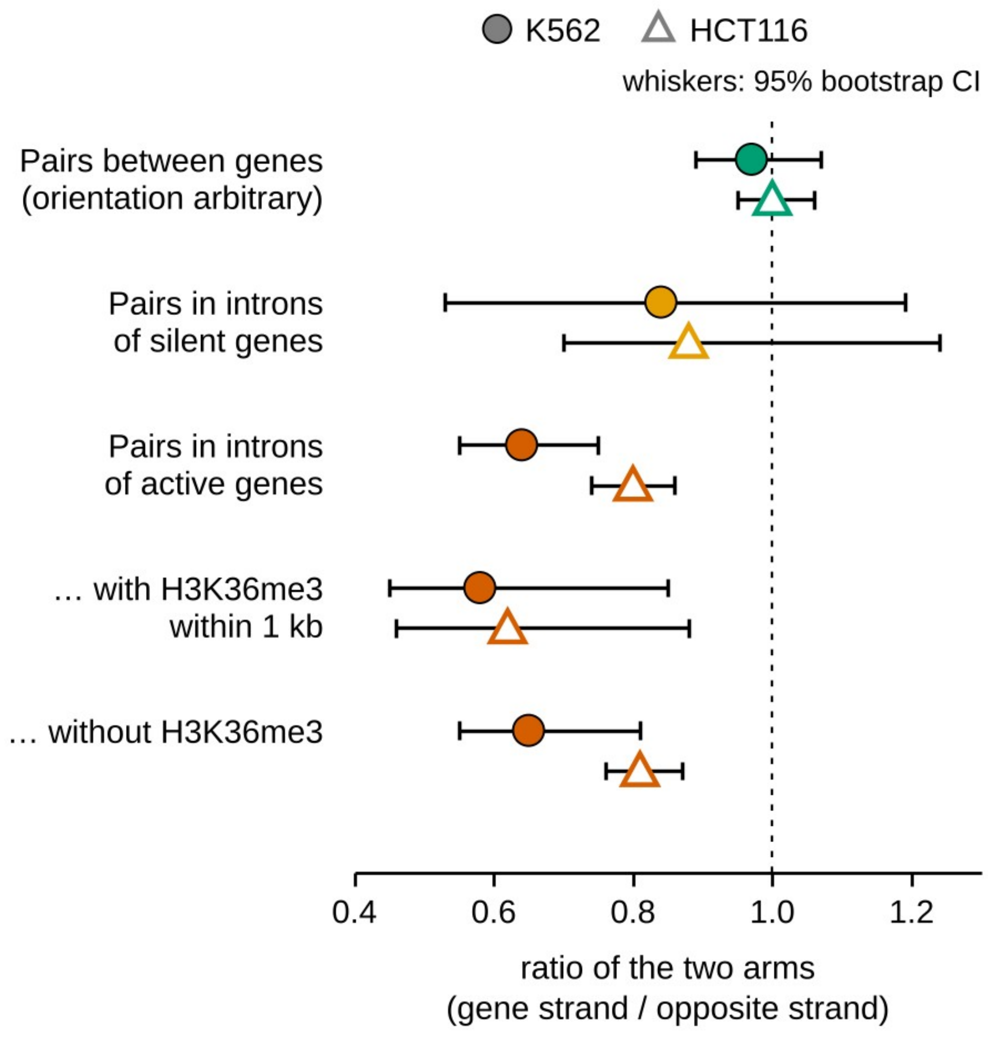


**Figure S8. Inside genes, initiation on the gene's own strand is weaker.** Median ratio between the two arms of divergent start pairs, the arm on the host gene's strand divided by the opposite arm, for pairs between genes (orientation arbitrary) and for pairs inside introns of silent and active genes (1,611 and 2,489 pairs in active genes, 252 and 369 in silent genes, in K562 and HCT116); pairs in active genes are also split by the presence of an H3K36me3 peak within 1 kb; whiskers show 95% bootstrap confidence intervals of the median (1,000 resamples). Active versus silent genes: $p = 0.42$ (K562) and 0.68 (HCT116); with versus without H3K36me3: $p = 0.88$ and 0.07 (Mann–Whitney tests). Arm on the host's strand against the opposite arm: $p = 9 \times 10^{-9}$ (K562) and $2 \times 10^{-12}$ (HCT116) in active genes and $p = 0.14$ and 0.03 in silent genes, Wilcoxon signed-rank test. After matching each intronic pair with the pair between genes of the closest total reads (Methods), the deficit of the arm on the host's strand in active genes was 33% (95% CI 16–46%; $p = 1 \times 10^{-4}$) in K562 and 17% (8–26%; $p = 2 \times 10^{-7}$) in HCT116, and in silent genes 7% (−65% to 55%; p = 0.49) and 7% (−33% to 31%; p = 0.27) (confidence intervals from resampling whole host genes; Wilcoxon signed-rank test on the matched pairs). Correlation of the asymmetry with the strength of the host promoter: Spearman ρ = −0.01 (p = 0.83) and −0.06 (p = 0.005).

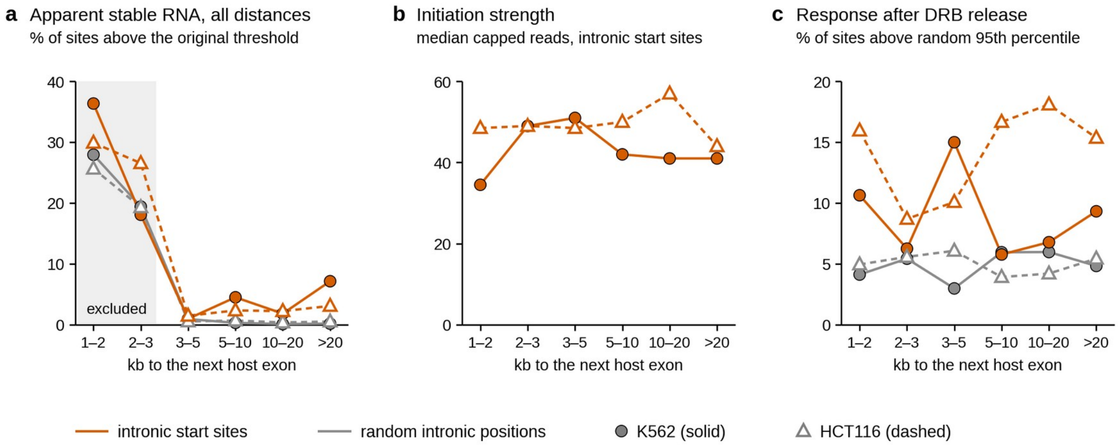


**Figure S9. Distance to the next exon of the host gene.** (a) Share of sites with an apparent rise in stable RNA under the original measure (total RNA-seq 0–3 kb after against 0.2–3.2 kb before the site, threshold from all random positions), by distance to the next exon of the host's reference transcript. Within 3 kb the window after the site contains that exon, and random intronic positions show the same apparent rise as intronic start sites; stable RNA was therefore measured only for sites at least 3 kb upstream of the next host exon (shaded bins excluded). (b) Initiation strength of intronic start sites in active hosts by the same distance (Spearman ρ = −0.02, p = 0.45, in K562; −0.04, p = 0.04, in HCT116). (c) Share of sites with a nascent RNA response 10 minutes after DRB release (BruDRB-seq in K562, DRB/TTchem-seq in HCT116), above the 95th percentile of random positions.

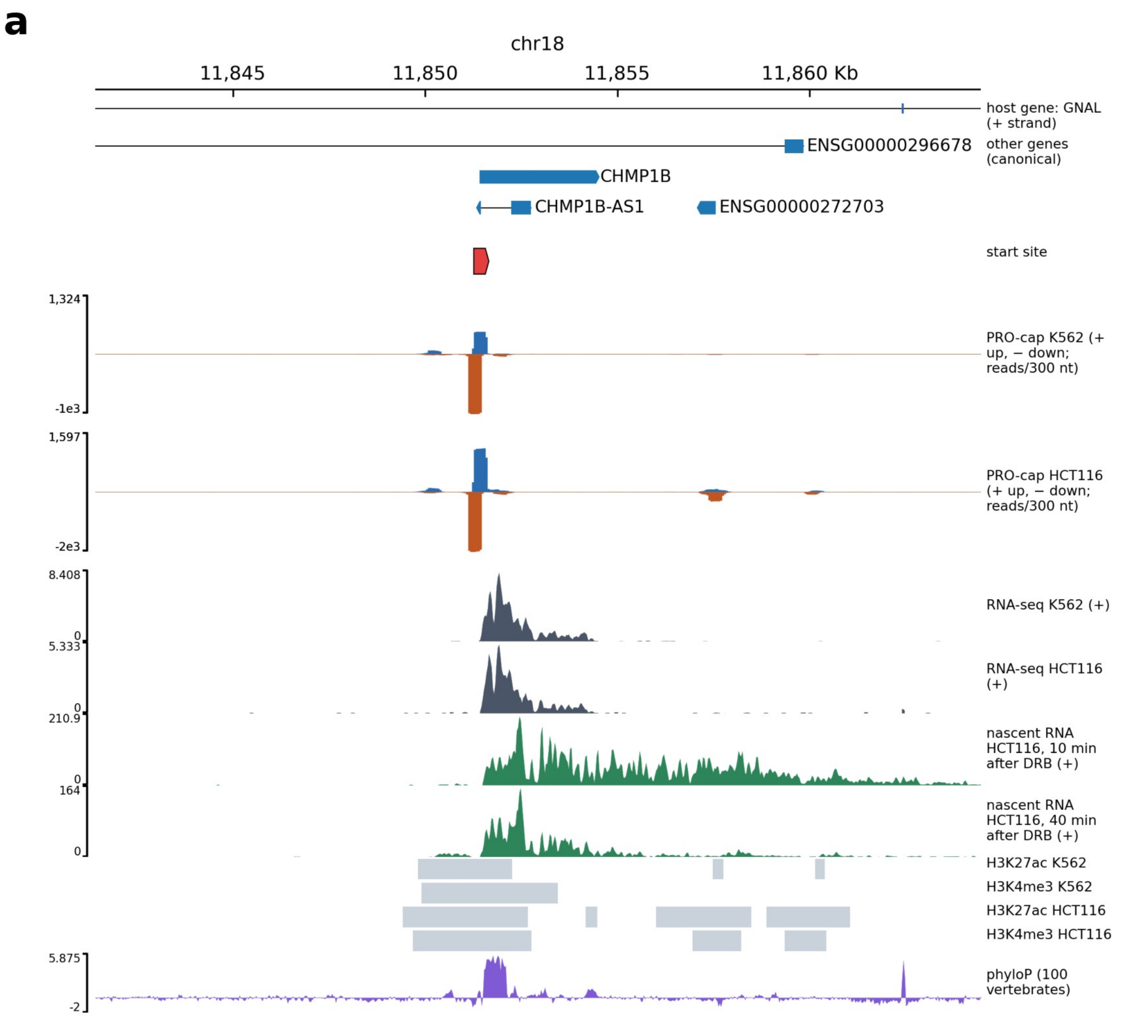
a
chr18
11,845
11,850
11,855
11,860 Kb
host gene: GNAL (+ strand)
ENSG00000296678
other genes (canonical)
CHMP1B
CHMP1B-AS1
ENSG00000272703
start site
1,324
-1e3
PRO-cap K562 (+ up, − down; reads/300 nt)
1,597
-2e3
PRO-cap HCT116 (+ up, − down; reads/300 nt)
8.408
0
RNA-seq K562 (+)
5.333
0
RNA-seq HCT116 (+)
210.9
0
nascent RNA HCT116, 10 min after DRB (+)
164
0
nascent RNA HCT116, 40 min after DRB (+)
H3K27ac K562
H3K4me3 K562
H3K27ac HCT116
H3K4me3 HCT116
5.875
-2
phyloP (100 vertebrates)

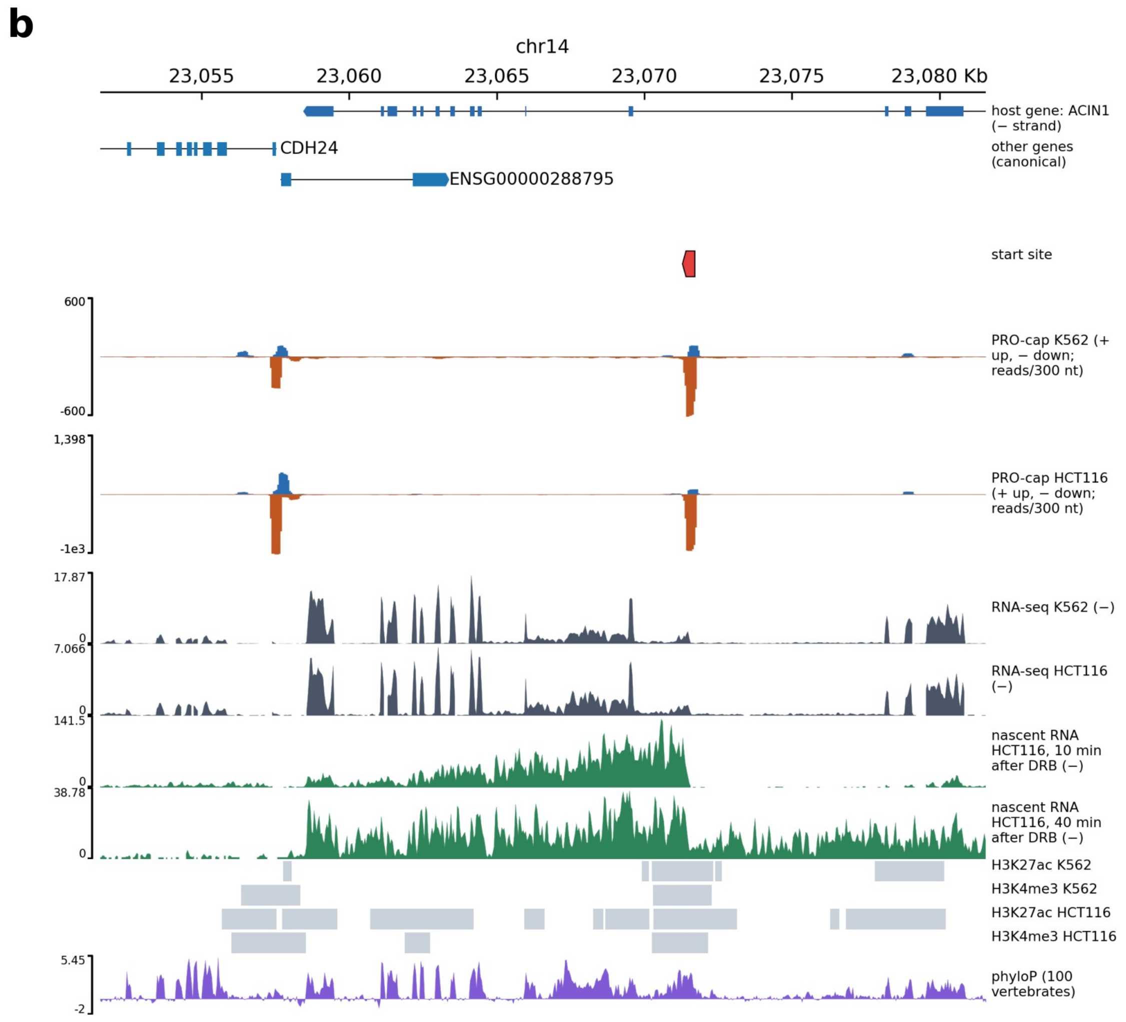


**Figure S10. Reference loci: a nested gene and an internal promoter.** Tracks as in Fig. 6. (a) CHMP1B, a known protein-coding gene nested in GNAL: a strong divergent start in both cell lines (its antisense partner is CHMP1B-AS1), stable RNA across the gene, transcription that continues and high conservation. (b) The internal promoter of ACIN1 (red mark, on the minus strand like ACIN1), an annotated alternative start of the gene: its RNA joins the host's exons, and nascent RNA runs about 13 kb in 10 minutes.

#### Tables

##### Table S1. All intronic start sites

Additional file 2, an Excel file (one sheet with the data, one describing the columns), lists the 5,801 intronic start sites of both cell lines, one row per site and cell line, with every measure used in the paper; stable RNA measures are given only for sites at least 3 kb upstream of the next exon of the host gene. Some columns apply to one cell line only (the DRB class to HCT116, the MTREX knockout to K562), and the measures per initiation event were computed only for unannotated sites, as in Fig. 5; other cells are empty.

| Column | Meaning |
|---|---|
| cell_line, site_id | cell line and site identifier |

| Column | Meaning |
| --- | --- |
| chrom, strand, position | genomic position of the start site (GRCh38, 0-based), on the host gene's strand |
| host_gene, host_active | host gene; whether its start in use carries at least five capped reads |
| capped_start_confirmed | whether the site passed the cap test |
| procap_reads | capped PRO-cap reads within 100 nt of the site |
| chromatin_type | promoter-like, enhancer-like or unmarked |
| H3K4me3, H3K27ac, H3K4me1 | whether a peak of the mark lies within 1 kb |
| phylop_mean | mean phyloP score over 201 nt centred on the site |
| stable_rna_rise_log2 | log2 of RNA-seq coverage 0–3 kb after the site over 0.2–3.2 kb before it |
| stable_rna_per_initiation | excess of RNA-seq coverage after the site over before it, per PRO-cap read |
| pas_per_kb_transcribed_strand, pas_per_kb_opposite_strand | polyadenylation signals per kb in the first kilobase on each strand |
| splice_donor_before_first_pas | whether the first splice-donor motif precedes the first polyadenylation signal |
| repeat_class, repeat_element, repeat_orientation | RepeatMasker element covering the start nucleotide and its orientation relative to transcription |
| drb_class_10min | single-site class in HCT116: productive, short, no response or unresolved |
| drb_far_per_initiation | excess of nascent RNA 2–8 kb after the site 10 minutes after DRB release, per PRO-cap read |
| mtrex_knockout_change | change of the stable RNA rise in MTREX knockout relative to control cells |
| shared_with_other_cell_line | whether a start site on the same strand lies within 200 nt in the other cell line |
| distance_to_next_host_exon | distance (nt) from the site to the next exon of the host's reference transcript in the direction of transcription; the stable RNA columns are empty below 3,000 |

**Table S2. Data sources**

| Data | Cell line | Accession | Used for |
|---|---|---|---|
| PRO-cap, capped RNA (signal, two replicates) | K562 | ENCSR261KBX | start sites, cap test, divergent pairs |
| PRO-cap, uncapped RNA (background) | K562 | ENCSR114HGS | cap test |
| PRO-cap peaks (PINTS), unidirectional and bidirectional | K562 | ENCFF636REV, ENCFF819CPA | intronic start sites, divergent pairs |
| PRO-cap and PINTS peaks, untreated CTCF degron line | HCT116 | ENCSR165QKR | start sites, cap test (main) |
| PRO-cap and PINTS peaks, untreated RAD21 degron line | HCT116 | ENCSR635SXX | reproducibility |
| Total RNA-seq (signal and alignments) | K562 | ENCSR792OIJ | stable RNA, splicing, expression |
| Total RNA-seq (signal and alignments) | HCT116 | ENCSR698RPL | stable RNA, splicing, expression |
| H3K4me3, H3K27ac, H3K4me1 ChIP-seq peaks | K562 | ENCSR668LDD, ENCSR000AKP, ENCSR000AKS | chromatin type |
| H3K36me3 ChIP-seq peaks | K562 | ENCSR000AKR | strand asymmetry |
| H3K4me3, H3K27ac, H3K4me1, H3K36me3 ChIP-seq peaks | HCT116 | ENCSR333OPW, ENCSR661KMA, ENCSR161MXP, ENCSR091QXP | chromatin type, strand asymmetry |
| RNA-seq, MTREX knockout and control | K562 | ENCSR300UBK, ENCSR271ZAB | exosome |
| RNA-seq, EXOSC10 knockout and control | K562 | ENCSR299PRA, ENCSR854OFV | exosome |
| RNA-seq, EXOSC5 knockout and control | K562 | ENCSR406SEH, ENCSR341TTW | exosome (failed check) |
| BruDRB-seq 10 min and Bru-seq | K562 | GSE55534 (SRR1182352–SRR1182355) | nascent RNA after DRB |
| DRB/TTchem-seq 10, 20, 30, 40 min, wild type | HCT116 | GSE148844 (SRR11563139–SRR11563142, SRR11563147–SRR11563150) | nascent RNA after DRB, single-site classes |
| Genome GRCh38 primary assembly | – | GENCODE | all analyses |

| Data | Cell line | Accession | Used for |
|---|---|---|---|
| and GENCODE release 47 annotation | | | |
| PolyASite 2.0 clusters | – | atlas.clusters.2.0.GRCh38.96 | polyadenylation sites |
| phyloP, 100 vertebrates | – | UCSC hg38.phyloP100way | conservation |
| RepeatMasker | – | UCSC hg38 rmsk | transposable elements |
| Median expression per tissue | – | GTEx v8 (RNASeQCv1.1.9) | broadly expressed genes |
